# An atlas of amyloid aggregation: the impact of substitutions, insertions, deletions and truncations on amyloid beta fibril nucleation

**DOI:** 10.1101/2022.01.18.476804

**Authors:** Mireia Seuma, Ben Lehner, Benedetta Bolognesi

## Abstract

Multiplexed assays of variant effects (MAVEs) guide clinical variant interpretation and reveal disease mechanisms. To date, MAVEs have focussed on a single mutation type - amino acid (AA) substitutions - despite the diversity of coding variants that cause disease. Here we use Deep Indel Mutagenesis (DIM) to generate the first comprehensive atlas of diverse variant effects for a disease protein, the amyloid beta (Aß) peptide that aggregates in Alzheimer’s disease (AD) and is mutated in familial AD (fAD). The atlas identifies known fAD mutations and reveals many variants beyond substitutions accelerate Aß aggregation and are likely to be pathogenic. Truncations, substitutions, insertions, single- and internal multi-AA deletions differ in their propensity to enhance or impair aggregation, but likely pathogenic variants from all classes are highly enriched in the polar N-terminus of Aß. This first comparative atlas highlights the importance of including diverse mutation types in MAVEs and provides important mechanistic insights into amyloid nucleation.

## Introduction

Amyloid fibrils are the hallmarks of more than 50 human diseases, including Alzheimer’s disease (AD), Parkinson’s disease, frontotemporal dementia, amyotrophic lateral sclerosis and systemic amyloidoses^1^. Mutations in the proteins that aggregate in the common forms of neurodegeneration also cause rare familial neurodegenerative diseases. For example, amyloid plaques of the amyloid beta (Aß) peptide are a pathological hallmark of AD and specific dominant mutations in Aß also cause familial Alzheimer’s disease (fAD)^2,3^.

The structures of many amyloid fibrils have now been determined^4^ including those of Aß fibrils extracted post-mortem from AD patient brains^5^. In these fibrils, the peptide adopts an S-shaped fold from residue 19 to 42, with the aliphatic C-terminus 29-42 packed as the inner core of the amyloid fibril. A more exposed N-terminal arm connects this to the first part of the peptide which remains unstructured in mature fibrils (residues 1-9 in sporadic and 1-11 in fAD). Despite these high-resolution structures, the mechanism by which fibrils form in the first place - the nucleation reaction - is still poorly understood, even though this is the fundamental process that needs to be understood and targeted to prevent amyloid diseases^6,7^. Moreover, we have only a superficial understanding of how specific mutations accelerate the process of amyloid nucleation to cause familial diseases. In Aß, several of the known fAD mutations are at residues outside the structured amyloid core^8^. Amongst other consequences, this makes the clinical interpretation of genetic variants challenging, with the vast majority of mutations identified in aggregating proteins classified as variants of uncertain significance (VUSs)^9^.

Multiplexed assays of variant effects (MAVEs)^10^ use cell-based or *in vitro* selection assays to build comprehensive atlases of variant effects (AVEs)^11^ to guide the clinical interpretation of VUSs^11^. This approach, which is also called deep mutational scanning (DMS), uses massively parallel DNA synthesis, selection and deep sequencing to quantify the relative activities of variants in a functional assay^12^. Applied to disease genes, DMS can also reveal disease mechanisms and it can be used to genetically-validate the relevance of cellular and *in vitro* disease models to human disease^13^. For example, we recently adopted a cell-based assay^14^ to allow massively parallel quantification of variant effects on protein aggregation. Measuring the effects of single nucleotide changes in Aß revealed that the assay both accurately quantifies the rate of amyloid fibril nucleation and that it identifies all of the dominant substitutions known to cause fAD^15^.

To date MAVE experiments^11^ have focussed on a single type of mutation - amino acid (AA) substitutions - and have largely ignored additional forms of genetic variation. Insertions and deletions (indels), in particular, are an abundant and important class of genetic variation in protein coding regions known to cause many human genetic diseases^16,17^, with small indels (<21 bp) causing approximately 24% of Mendelian diseases^18,19^. Indels are a fundamentally different perturbation to a protein sequence to substitutions: whereas substitutions only alter AA side chains, indels are backbone mutations that change the length of the polypeptide chain and so may be expected to have more severe effects^20^. However, despite their importance, there has been very little systematic quantification of the effects of indels in proteins^21–23^, particularly in disease genes, and many computational methods for predicting variant effects simply ignore them^24^. To our knowledge, a systematic comparison of the effects of AA substitutions, insertions and deletions is lacking for any human disease gene.

Here we address this fundamental shortcoming in human genetics by providing the first comprehensive comparison of the effects of substitutions, insertions and deletions in a human disease gene.

The resulting AVE quantifies the effects of diverse sequence changes on the aggregation of Aß and is the first dataset that can be used to guide the clinical interpretation of different types of mutation in a human disease gene. It reveals that many mutations beyond substitutions accelerate the aggregation of Aß and so are likely to be pathological. The atlas identifies the two deletions known to cause fAD, but reveals that they are only two of the many insertions and deletions that are likely to be pathological. The atlas also provides fundamental mechanistic insight into the process of amyloid nucleation, illustrating the power of deep indel mutagenesis (DIM) to illuminate sequence-to-activity relationships.

## Results

### Deep Indel Mutagenesis of amyloid beta

To quantify and contrast the effects of diverse genetic variants on the aggregation of the 42 AA form of Aß (Aß42), which is the most abundant component of amyloid plaques in AD, we performed Deep Indel Mutagenesis (DIM) by synthesizing a library containing all possible single AA substitutions (n=798), all possible single AA insertions (n=780), all single AA deletions (n=37), all internal multi-AA deletions ranging in length from 2-39 AA (n=731), and all progressive truncations from the N-terminus, C-terminus or both, removing 2-39 AA (n=817, Fig. 1a).

**Figure 1.**
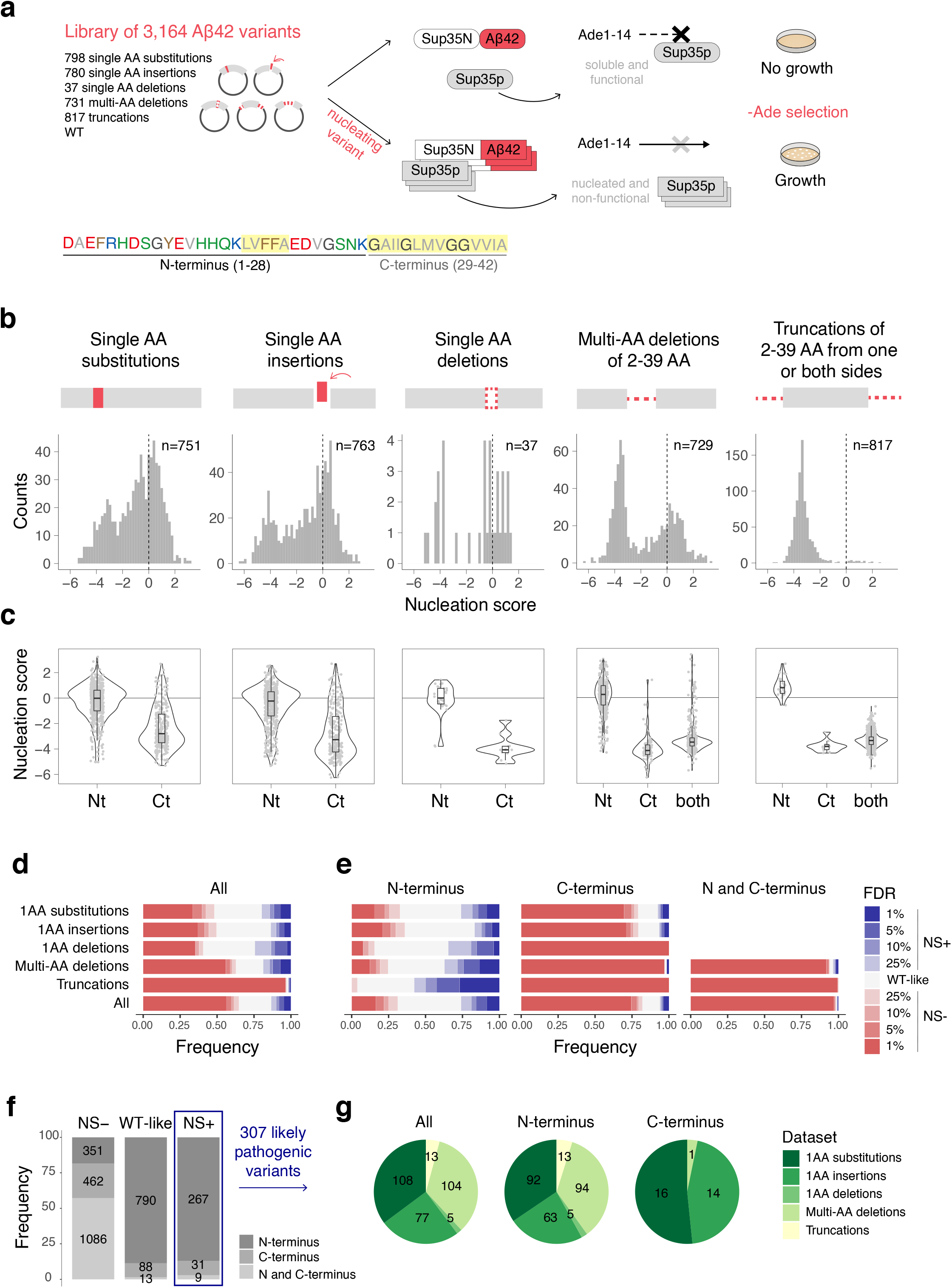
Deep indel mutagenesis of Aß. **a** Aß coding sequence colored by AA class (red: negative, blue: positive, green: polar, gray: aliphatic, brown: aromatic, dark gray: glycine) and schematics of the *in vivo* selection assay. Aß, fused to sup35N, seeds aggregation of sup35p causing a read-through of a premature stop codon in the *ade2* reporter gene allowing growth in medium lacking adenine. **b** Distribution of nucleation scores for each class of mutations. Dashed lines indicate WT nucleation score (0). **c** Distributions of nucleation scores for mutations in different regions: N-terminus (AA 1-28), C-terminus (AA 29-42) or both (AA 1-42). **d,e** Frequency of variants increasing or decreasing nucleation at different FDRs for the full peptide (**d**) and for each peptide region (**e**). **f** Frequency and total counts of each mutation type for variants increasing (NS+), decreasing (NS-) or having no effect (WT-like) at FDR=0.1. **g** Number and type of variants increasing nucleation (NS+) for each peptide region.

We quantified the effects of these different classes of variants in a cell-based selection assay where the aggregation of Aß nucleates the aggregation of an endogenous protein, a process required for growth in selective conditions (Fig. 1a)^14,15^. After selection, the enrichment of each variant in the library was quantified by deep sequencing^15,25^. The resulting enrichment scores are reproducible between replicates (Supplementary Fig. 1a) and correlate well with previous measurements (R=0.82, Supplementary Fig. 1b) as well as with the effects of variants quantified individually (R=0.89, Supplementary Fig. 1c). In addition, and as previously reported^15^, the enrichment scores correlate linearly with the *in vitro* measured kinetic rate constants of Aß amyloid fibril nucleation (R=0.96, Supplementary Fig. 1d)^15,26^, so we refer to them as “nucleation scores”.

### Contrasting the impact of substitutions, deletions and insertions in a disease gene

The resulting dataset provides the first opportunity to comprehensively compare the effects of different types of mutation - substitutions, insertions, deletions and truncations - in a human disease gene. Focussing on single AA changes, the most frequent mutational effect is reduced aggregation, with 43% of substitutions, 44% of insertions, and 37% of deletions having lower nucleation scores (NS) than wild-type (WT) Aß (false discovery rate, FDR=0.1, NS-variants, Fig. 1b,d). The effects of multi-AA deletions are stronger, with 60% of internal multi-AA deletions and 97% of multi-AA truncations from one or both ends reducing nucleation (FDR=0.1, Fig. 1b,d).

### Many variants beyond substitutions accelerate Aß aggregation

Variants in Aß identified in families with fAD accelerate Aß aggregation, consistent with a gain-of-function mechanism^15,27^. Unlike computational methods to predict aggregation or variant effects, the experimental nucleation scores accurately classify fAD variants (Supplementary Fig. 1e). In total, there are 307 variants in our library (10%) that accelerate Aß aggregation (FDR=0.1, NS+ variants): 108 substitutions, 77 insertions, 5 single AA deletions, 104 internal multi-AA deletions and 13 truncations (Fig. 1f,g and Supplementary Table 1). There are thus many variants beyond substitutions that accelerate the aggregation of Aß.

### All types of variant that promote aggregation are strongly enriched in the N-terminus

The primary sequence of Aß consists of an N-terminal region enriched in charged and polar residues (AA 1-28, two thirds of the peptide) and a C-terminal region composed entirely of aliphatic residues and glycines (AA 29-42, one third of the peptide) (Fig. 1a).

For all classes of mutation, variants that reduce nucleation are strongly enriched in the aliphatic C-terminus of Aß: 60% of substitutions, 54% of insertions, 78% of single AA deletions, 85% of the internal multi-AA deletions and all truncations that reduce nucleation occur in this hydrophobic region (FDR=0.1; Fig. 1c,f and Supplementary Fig. 1f). Indeed for all mutation types, the majority of variants in this region impair nucleation: 76% of substitutions, 76% of insertions, all single AA deletions, 94% of internal multi-AA deletions and all truncations (Fig. 1e).

In contrast, variants that accelerate nucleation are strongly enriched in the polar N-terminus. In total, 87% of variants that accelerate nucleation (267/307, FDR=0.1, NS+ variants) are located in the polar N-terminus (Fig. 1f). This contrasts to just 18% of variants that reduce nucleation (Fig. 1f). This strong enrichment is true for all mutation types: 85% of substitutions, 82% of insertions, 90% of multi-AA deletions, all single AA deletions, and all truncations that accelerate nucleation occur in the N-terminus (Fig. 1c,f,g). Very few variants in the aliphatic C-terminus increase nucleation: only 16 substitutions (6%) and 14 insertions (5%), while none of the single AA deletions do so. Similarly, no C-terminal truncations accelerate nucleation and only one internal multi-AA deletion in the C-terminus does so (Fig. 1e-g).

### Mutation classes differ in their propensity to promote or prevent amyloid nucleation

The different classes of mutation do, however, vary in how likely they are to increase or decrease nucleation when they occur in the same region. The type of mutation most likely to accelerate nucleation is N-terminal truncations, with 50% increasing nucleation and no N-terminal truncation reducing nucleation (FDR=0.1, Fig. 1e). More internal multi-AA deletions in the N-terminus increase than decrease nucleation (28% vs. 19%), as do more single AA deletions (19% vs 11%). In contrast, single AA substitutions in the N-terminus are more likely to decrease (26%) than increase (18%) nucleation, as are insertions (30% decrease and 12% increase) (FDR=0.1, Fig. 1e).

In summary, the DIM data reveals that there are many mutations beyond single AA substitutions that accelerate Aß aggregation and so are potentially pathogenic (Supplementary Table 1). Moreover, they show that, for all mutation types, the vast majority of variants that accelerate nucleation are located in the polar and charged N-terminal region of Aß. However, the different classes of mutation have very different distributions of mutational effects in the N-terminus: whereas single AA substitutions and insertions in the N-terminus are more likely to decrease nucleation than increase it, the opposite is true for single AA deletions, internal multi-AA deletions and N-terminal truncations: these mutation classes more often enhance nucleation than impair it, suggesting they are particularly likely to be pathogenic if they occur.

### AA preferences in the N-terminus: polar, positive, small and P residues promote nucleation

Considering all positions, the effect of substituting in an AA is moderately correlated to the effect of inserting the same AA before or after the same position (R=0.49 and R=0.51, respectively, Fig. 2e). This relationship is, however, partly driven by the distinct impact of mutations at the N and the C-terminus (Supplementary Fig. 3). Thus, although the consequences of insertions and substitutions are related, they are also clearly distinct, as is also revealed by comparing their effects at each individual residue (Supplementary Fig. 5a) and their average effects across all residues (Supplementary Fig. 4a).

**Figure 2.**
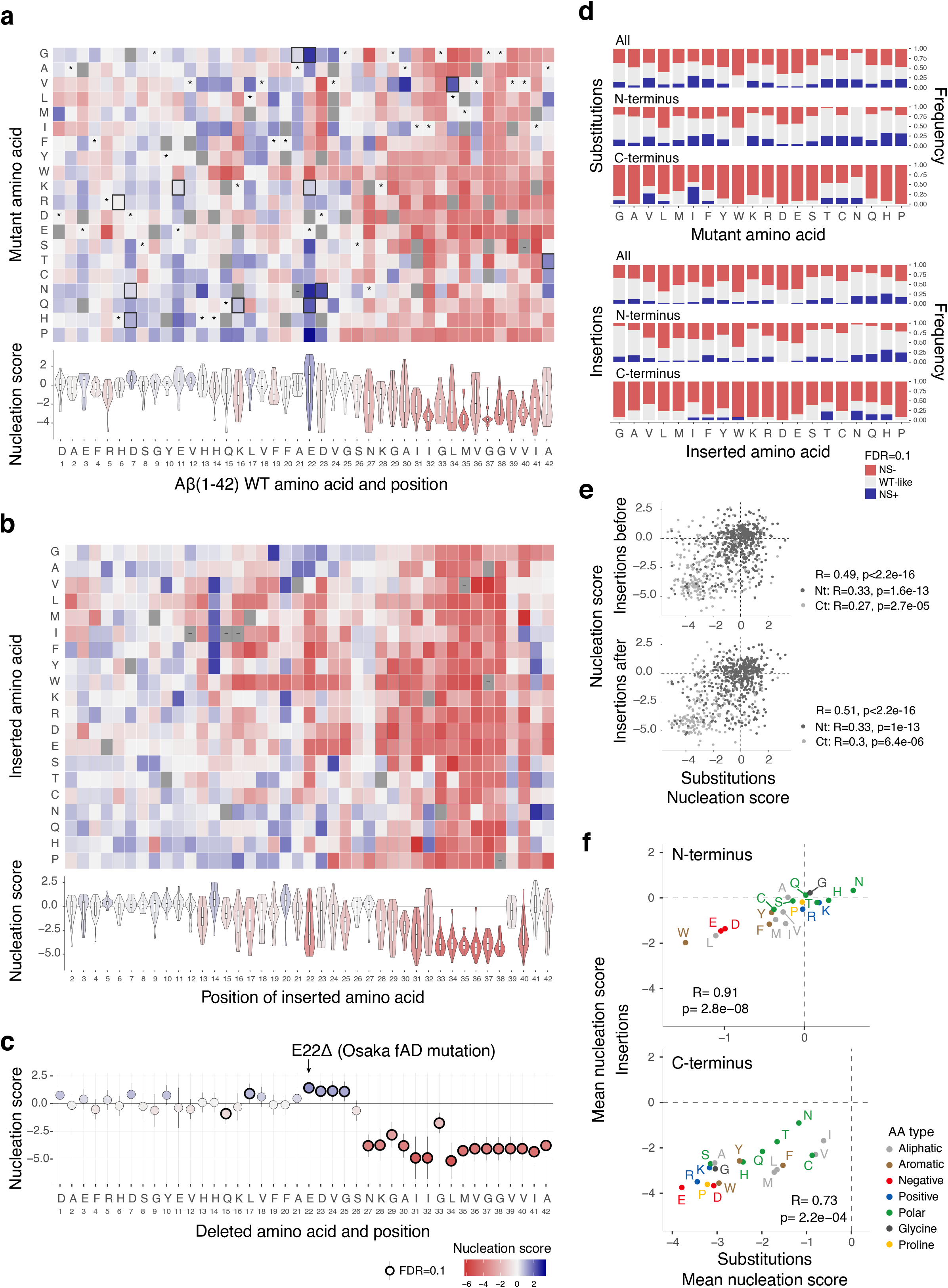
Single AA variant atlases. **a** Heatmap of nucleation scores for single AA substitutions. The WT AA and position are indicated in the x-axis and the mutant AA is indicated in the y-axis. Variants not present are indicated in gray, synonymous mutants with ‘*’ and fAD mutants with a black box. Non-nucleating variants (with no NS, see methods) are indicated with ‘-’. The distribution of nucleation scores for each position is summarized in the violin plots below the heatmap. **b** Heatmap of nucleation scores for single AA insertions at each position. **c** Effect of single AA deletions. The horizontal line indicates the WT nucleation score (0). Vertical error bars indicate 95% confidence interval of the mean. **d** Frequency of AA increasing or decreasing nucleation (FDR=0.1) upon substitutions (top) or insertion (bottom) for each peptide region. **e** Correlation of nucleation scores for substitutions to each AA at each position and insertions of the same AA before (top) or after (bottom) that position. Color indicates peptide region (N and C-terminus). **f** Correlation of average nucleation scores for each AA, for insertions and substitutions at the N-terminus (top) and at the C-terminus (bottom). Color indicates AA type. Pearson correlation coefficients are indicated in (**e**) and (**f**).

Considering the N and C-terminal regions separately, the average effects of inserting or substituting in AAs in any positions are strongly related (R=0.91 and R=0.73 for residues 1-28 and 29-42, respectively, Fig. 2f and Supplementary Fig. 4b,c). Both substituting and inserting polar residues (especially, N,H,T,Q) into the N-terminus frequently promotes aggregation, as does adding positively charged residues (K,R). Interestingly, substituting in or inserting G or A into the N-terminus also frequently increases nucleation as does adding a P, particularly in the second half of the N-terminus (Fig. 2a,b,d and Supplementary Figs. 2a,b and 6-8). Since P residues are unlikely to be tolerated in the core of structured fibrils, their effect in promoting nucleation may be via changes in the ensemble of soluble Aß^28^, rather than due to changes in the fibril transition state. For example, adding P might impair the formation of a transient secondary structure that - in the WT ensemble - acts to prevent nucleation.

Overall, these enrichments for polar, positively charged, small and P residues are very different to the sequence preferences used by computational methods to predict protein aggregation^29–32^, and these methods indeed perform very poorly for predicting the effects of mutations in the N-terminus of Aß (Supplementary Fig. 9).

### AA preferences in the N-terminus: increased hydrophobicity and negatively charged residues reduce aggregation

Also inconsistent with the expectations of predictive methods, the substitutions and insertions in the N-terminus that most often reduce aggregation are additions of hydrophobic (W,L,F,M,I,Y,V) and negatively charged (D,E) AAs (Fig. 2a,b,d and Supplementary Fig. 2a,b). Consistent with this, individually deleting negatively charged residues and L17 from the N-terminus often increases nucleation (Fig. 2c), as does substituting away from these same AAs (Supplementary Fig. 2c).

However, there are many exceptions to these general trends, highlighting the importance of generating the full mutational matrix. For example, W insertions and substitutions to W mostly promote aggregation in residues 1-12 but nearly always impair aggregation at positions 13-28 (Fig. 2a,b and Supplementary Fig. 2a,b). In addition, many substitutions to V and I in positions 13-20 strongly increase aggregation, as do many hydrophobic insertions after residue 13 between two histidines. At particular positions the impact of substitutions or insertions can also be quite distinct: for example substitutions of F19 and F20 rarely increase nucleation and only for mutations to hydrophobic AA, while many insertions between F19 and F20, increase nucleation, especially of charged and polar residues (Fig. 2a,b, and Supplementary Fig. 2a,b). At other residues the preferences are more similar: for example both substitutions to and insertions of G at residues 22 and 23 increase nucleation resulting in some of the fastest nucleating variants in the library (Fig. 2a,b and Supplementary Figs. 2a,b and 13), suggesting that increasing flexibility or reducing side chain volume in this region favors nucleation.

Thus, although simple rules can predict mutational effects to some extent, the comprehensive Aß data suggests full experimental datasets and new computational methods will be required for the clinical interpretation of variants in aggregating proteins.

In summary, the role of the N-terminal two thirds of Aß in promoting and preventing amyloid nucleation must be very different to that of the C-terminus that forms the hydrophobic core of Aß fibrils. To our knowledge, no existing mechanistic models can satisfactorily account for mutational effects in this region^33,34^. In mature Aß fibrils derived from patients^5^ and formed *in vitro*^5,35–39^, part or all of the N-terminus remains unstructured (Fig. 5 and Supplementary Fig. 14). Changes in aggregation caused by mutations in this region could be due to their effects on the ensemble of soluble Aß. Alternatively, the N-terminus could participate directly in the nucleation reaction, establishing interactions in the nucleation transition state.

### The Osaka mutation (ΔE22) is the fastest nucleating single AA deletion

To date, only one single AA deletion has been reported in families with fAD: deletion of residue E22, named the Osaka mutation after the city in which it was first identified^40^. Strikingly, our data shows that the Osaka mutation is the single AA deletion that most enhances the nucleation of Aß (Fig. 2c). However, an additional 4 single AA deletions promote nucleation (FDR=0.1), suggesting that they may also be pathogenic. All of these deletions are in the N-terminus of the peptide (Fig. 2c).

### The Uppsala mutation (Δ19-24) lies in a hotspot of internal multi-AA deletions that promote Aß nucleation

After we generated this dataset, the first internal multi-AA deletion in Aß that causes fAD was reported^41^. This deletion, referred to as the Uppsala mutation, removes AA 19-24. The Uppsala mutation strongly promotes nucleation in our dataset (Fig. 3a,b). However, the comprehensive DIM altas also reveals that there are an additional 103 internal multi-AA variants that promote Aß aggregation (FDR=0.1; Fig. 3a,b, Supplementary Fig. 10a and Supplementary Table 1). Strikingly, however, the Uppsala mutation is located in the center of a hotspot region where many different deletions accelerate aggregation (Fig. 3a-d). In total, 35 multi-AA deletions removing some or all of residues 17-27 increase the nucleation of Aß (FDR=0.1, Fig. 3d and Supplementary Fig. 10a). This suggests that there are potentially many more pathogenic deletions that remain to be discovered that remove residues in this central hotspot region, as well as additional pathogenic deletions throughout the N-terminus (Supplementary Table 1).

**Figure 3.**
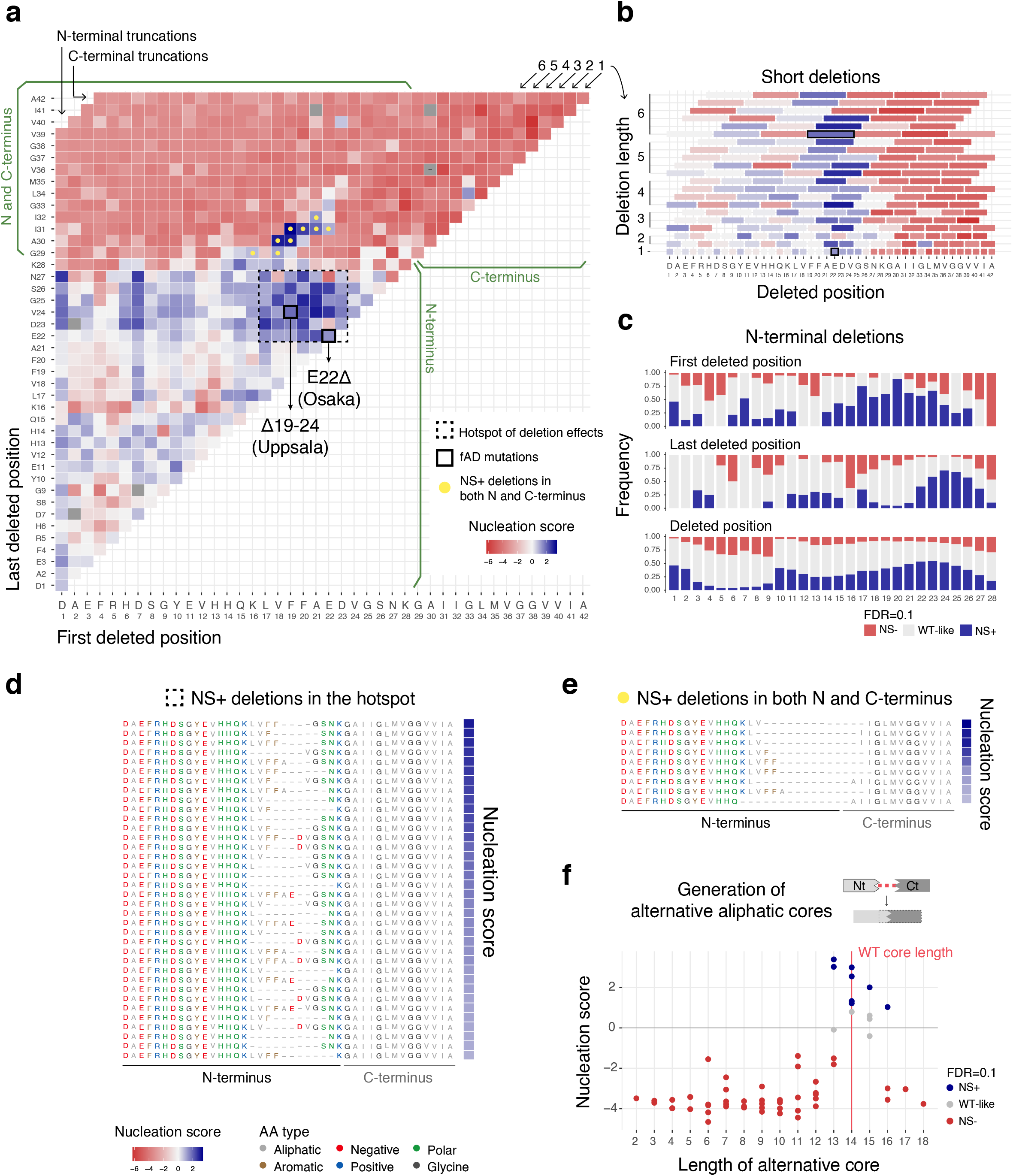
Internal multi-AA deletion atlas. **a** Matrix of nucleation scores for deletions. The dashed-line black square depicts the hotspot of deletion effects (consecutive deleted positions where NS+ frequency > ½ max(NS+ frequency, i.e. deletions starting at positions 17-23 and ending at positions 22-27) and the yellow dots indicate deletions removing residues in both the N and C-terminus that increase NS (see (**e**)). Variants not present are represented in gray and non-nucleating variants (with no NS, see methods) are indicated with ‘-’. **b** Effect on nucleation of deletions of 1-6AA length. The WT AA and position are indicated in the x-axis. The black squares indicate fAD variants: Osaka (E22Δ) and Uppsala (Δ19-24). Color code as in (**a**). **c** Frequency of variants increasing nucleation (NS+), decreasing nucleation (NS-) or with no difference from WT (WT-like) at FDR=0.1, for sequences with a specific first deleted position (i.e. each column in the matrix), last deleted position (i.e. each row in the matrix) or missing a specific residue, at the N-terminus (AA 1-28). **d** AA sequence for variants inside the hotspot of deletion effects with significantly increased NS (FDR=0.1). **e** AA sequence for variants with internal multi-AA deletions removing residues from both the N and C-terminus, with significantly increased NS (FDR=0.1). AA coloured by AA class. Color code as in (**d**). **f** Nucleation scores of variants with putative alternative aliphatic cores of different lengths. The horizontal line indicates the WT nucleation score (0). Vertical red line indicates WT core length (14AA). Variants displaying alternative cores were defined as those internal multi-AA deletions removing residues in both the N and C-terminus that replace part of the C-terminus with exclusively aliphatic residues (n=64).

The multi-AA deletion hotspot is centered on the negatively charged residues E22 and D23 (Fig. 3a-d and Supplementary Fig. 10a). Many substitutions at these two positions also accelerate nucleation (Fig. 2a) as does the individual deletion at position E22 (Osaka mutation, Fig. 2c). However, not all internal multi-AA deletions that remove E22 or D23 increase aggregation, with deletions starting from positions 4,5,12 and 13 that remove E22 or D23 failing to accelerate nucleation (Fig. 3a). In these cases, a negatively charged residue is relocated to the immediate proximity - one or two residues away - of the core (AA 29-42, Fig. 3a and Supplementary Fig. 10b), where they likely compensate for the loss of negative charge.

The importance of charge in mediating the effects of multi-AA deletions is also suggested by a cluster of deletions in the first 15 residues of Aß that accelerate nucleation (Fig. 3a and Supplementary Fig. 10a). This region contains four of the negatively charged residues in Aß (D1, E3, D7 and E11) with many substitutions of these residues also accelerating aggregation (Fig. 2a). The matrix of internal multi-AA deletions further reveals that deletions that remove D1 have higher NSs than those that keep it; the same is true for D7 (Fig. 3a and Supplementary Fig. 10a). This segment is unstructured in nearly all mature Aß fibril polymorphs^35–39^, including those in AD brains^5^ (Fig. 5 and Supplementary Fig. 14), yet diverse types of mutation in this region strongly increase aggregation.

### Mutations in the aliphatic core that accelerate aggregation

The vast majority of mutations of any type within the aliphatic C-terminus (AA 29-42) of Aß strongly disrupt nucleation (Fig. 1c,e,f). Indeed all insertions in the 33-38 stretch disrupt nucleation, suggesting that this may constitute the inner core of the nucleation transition state (Fig. 2b and Supplementary Fig. 2b).

However, there are some variants in the C-terminus that increase nucleation: 16 substitutions, 14 insertions, one internal multi-AA deletion within the C-terminus and nine multi-AA deletions that involve C-terminal residues (FDR=0.1, Fig. 1f,g). The substitutions in the C-terminus that accelerate nucleation are enriched at A30 and A42. At position 42, mutations to L promote nucleation, as do changes to C,T and N (Fig. 2a and Supplementary Fig. 2a). Among these, only A42T is a known fAD variant (Supplementary Table 4). At position 34 and 36, 4 substitutions to alternative hydrophobic AAs promote nucleation, suggesting that the L and V side chains may not be optimal in the nucleation transition state. L34V is also a known fAD variant (Supplementary Table 4). Insertions that promote nucleation are also enriched at specific positions. Polar insertions at position 32, flanking G33, may favor a turn, and polar, aromatic and hydrophobic insertions at positions 39, 41 and 42 (Fig. 2b and Supplementary Fig. 2b) at the end of the core may be more easily accommodated by minor structural rearrangements.

The only deletion within the core that accelerates nucleation is the removal of G33 and L34, although the individual deletion of each residue disrupts nucleation (Fig. 3a and Supplementary Fig. 10a). It is possible that adjustments in the core can accommodate removal of these two residues by the formation of a similar structural polymorph. Finally, nine internal multi-AA deletions that bridge the N and C-terminus increase nucleation (FDR=0.1, Fig. 3a,e and Supplementary Fig. 10a). These deletions remove aliphatic core residues but replace them with a similar number of aliphatic residues from a more N-terminal segment of the peptide (Fig. 3a,e). It is likely that these internal multi-AA deletions are therefore creating alternative aliphatic cores that nucleate to form the same or similar structural polymorphs as full length Aß. We find that these alternative cores that increase nucleation have a specific range of core lengths, with the hydrophobic stretch spanning from 13 to 16AA, very similar to the 14 AA length in the WT peptide (Fig. 3e,f and Supplementary Fig. 11a).

### Positive charge promotes the nucleation of a minimal Aß core

The DIM dataset shows that progressively removing AAs from the N-terminus of Aß generates many peptides that aggregate faster than the full 42AA isoform, with 13/27 N-terminal truncations promoting nucleation (Fig. 4a,b and Supplementary Fig. 12). Such N-terminally truncated fragments of Aß have been detected in AD patients^42,43^ (Supplementary Table 2) and our data suggests that environmental triggers, infections or genomic alterations that increase their production are likely to accelerate Aß aggregation and so may be causally important in familial and sporadic AD.

**Figure 4.**
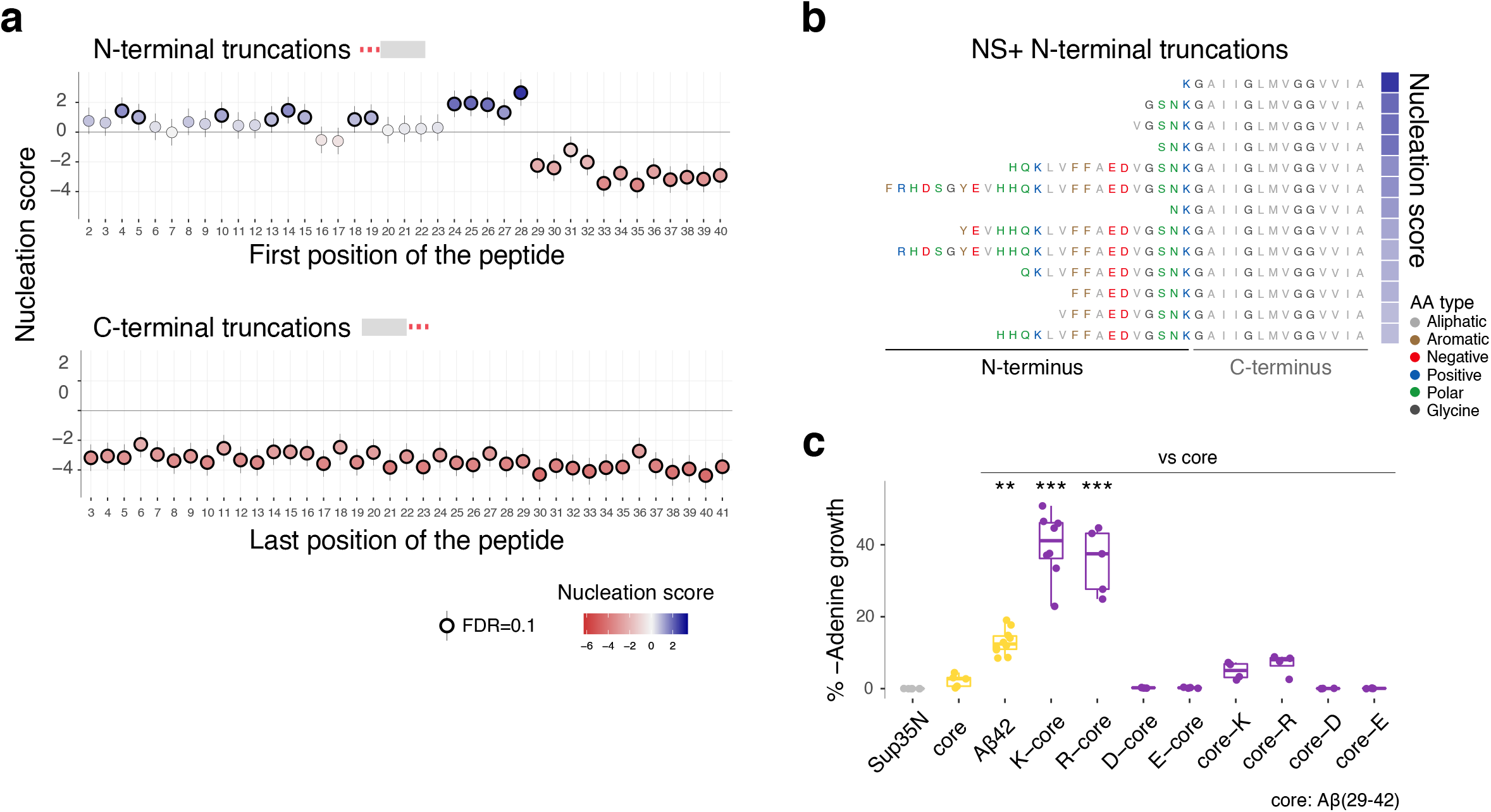
Positive charge accelerates nucleation of a minimal Aß core. **a** Effect of N-terminal (top) and C-terminal (bottom) truncations on nucleation. Vertical error bars indicate 95% confidence interval of the mean NS. **b** AA sequences of N-terminal truncations that increase nucleation at FDR=0.1. AA are coloured by class. **c** Effect of adding a positively or negatively charged residue at the N or C-terminus of the Aß core (AA 29-42). Nucleation quantified as percentage of colonies in medium lacking adenine vs. medium containing adenine. One-way ANOVA with Dunnett’s multiple comparisons test. * p<0.05, **p<0.01, *** p<0.001.

In contrast, all N-terminal truncations that remove at least one residue of the aliphatic core (AA 29-42) very strongly reduce aggregation, further highlighting the critical requirement for this region in nucleation (Fig. 4a and Supplementary Fig. 12). Strikingly, however, the aliphatic core alone nucleates very slowly (FDR=0.1, Fig. 4a). The addition of residue 28 to this minimal core dramatically accelerates nucleation, with the 15AA peptide consisting of residues 28-42 actually being the fastest nucleating N-terminally truncated form of Aß (Fig. 4a,b). This minimal Aß core nucleates faster than full-length Aß (Fig. 4a) and is too short to form the S-shaped amyloid fibrils polymorph observed in AD plaques^5^ and so likely adopts a smaller C-shaped polymorph with two main strands facing each other. The rapid nucleation of this 15AA peptide is particularly striking given the observation that all multi-AA deletions of more than 23AA prevent nucleation (Fig. 3a).

Residue 28 is a lysine and many of the other faster nucleating N-terminally truncated peptides also have positively charged residues at or close to their N-termini (Fig. 4b). Moreover, internal multi-AA deletions that remove K28 but that still nucleate often have a positively charged residue at the N-terminus of the core (Supplementary Figs. 10c and 11b). We therefore tested the hypothesis that it is the addition of a positively charged residue that accelerates nucleation of the minimal aliphatic core of Aß. Adding the positively charged residues K or R to the N-terminus of the Aß core (AA 29-42) strongly accelerated nucleation (Fig. 4c). In contrast, adding the negatively charged residues D or E did not (Fig. 4c). The addition of a single positively charged residue is therefore sufficient to dramatically accelerate the aggregation of the aliphatic core of Aß. It is possible that positive residues, but not negative ones, at position 28 engage in a salt bridge with the carboxyl group at the C-terminus of the peptide to promote nucleation^44^.

### Residue 42 is required for fast nucleation

In contrast to the effects of N-terminal truncations, removing even a single AA from the C-terminus of Aß strongly reduces nucleation (Fig. 4a and Supplementary Fig. 12). That A42 plays an important role in the nucleation of Aß is consistent with previous reports that Aß42 aggregates faster than Aß40^6^. However, position 42 does not need to be an A: multiple substitutions and multiple insertions before position 42 (Fig. 2a,b and Supplementary Fig. 2a,b) either do not disrupt nucleation or actually accelerate it. This suggests that the requirement for position 42 may therefore primarily be a steric one, for example to position a free carboxyl terminus in the nucleation transition state^44^.

## Discussion

We have presented here the first systematic comparison of the effects of substitutions, insertions and deletions in a human disease gene. The resulting dataset shows that the consequences of AA insertions, deletions and truncations are not trivial to predict from the effects of substitutions, highlighting the importance of including Deep Indel Mutagenesis (DIM) when constructing an atlas of variant effects (AVE)^11^ for the interpretation of clinical genetic variants.

The dataset provides a comprehensive AVE for Aß aggregation that can be used to guide the future clinical interpretation of variants as they are discovered. The atlas reveals that many variants beyond substitutions accelerate the aggregation of Aß and so are likely to be pathogenic. The identification of 307 variants that accelerate aggregation (Supplementary Fig. 13 and Supplementary Table 1) in this very short 42AA peptide highlights the potentially enormous diversity of disease-causing variants in the human genome. For example, the Aß AVE reveals that the Uppsala mutation (Δ19-24) is just one of many internal multi-AA deletions in a central hotspot region of Aß that accelerate aggregation; these additional deletions are also likely to be pathogenic, as are multiple additional single AA deletions in the N-terminus and many N-terminal truncations of the peptide.

The substitutions, insertions, deletions and truncations that accelerate nucleation are all strongly enriched in the polar N-terminal region of Aß (Fig. 5 and Supplementary Fig. 14). The different classes differ, however, in their distributions of mutational effects, with substitutions and insertions in the N-terminus more likely to impair rather than enhance aggregation but single and multi-AA deletions more likely to enhance rather than impair it. N-terminal truncations of Aß are particularly likely to accelerate nucleation, raising the intriguing possibility that increased production of N-terminally truncated forms of Aß triggered by environmental exposures, pathogens or genetics might be an important cause of familial and sporadic AD.

**Figure 5.**
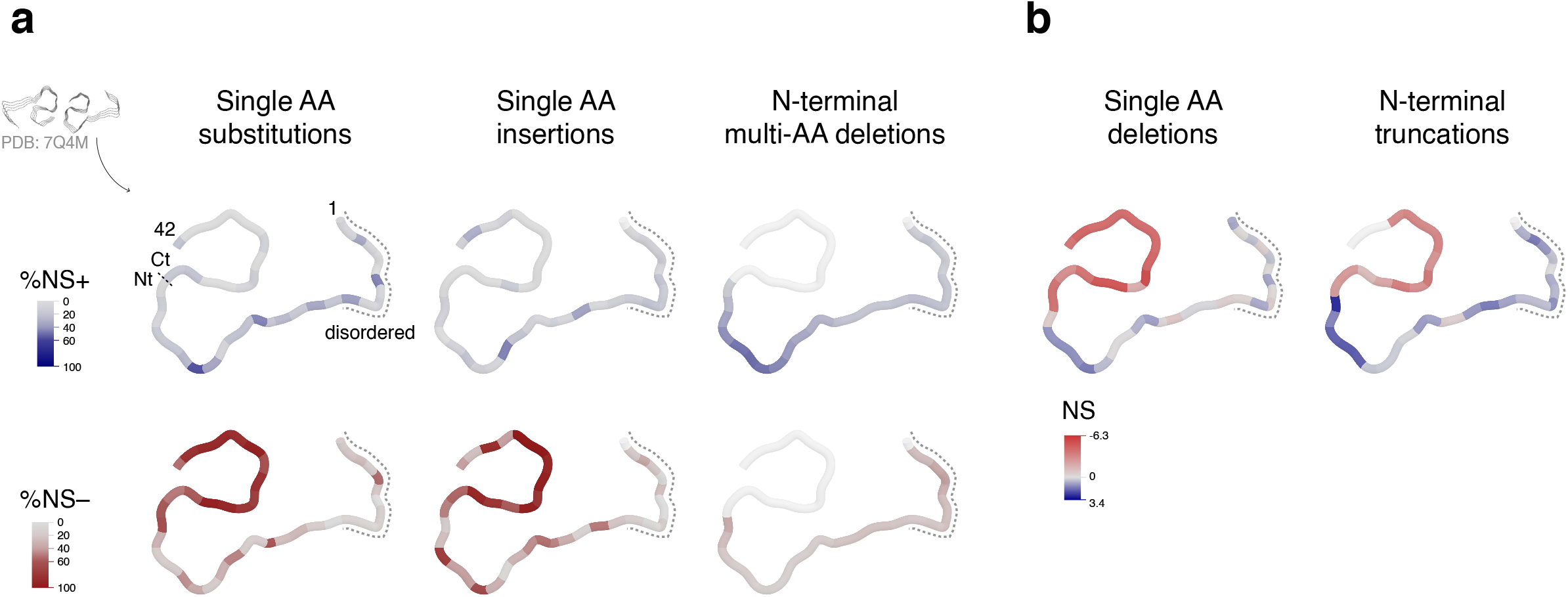
Mutational effects visualized on fAD Aß fibril structures. In fibrils extracted from the brains of fAD patients, Aß42 adopts an S-shaped structure at the C-terminus with an N-terminal arm linking to an unstructured region indicated by the dashed line (PDB: 7Q4M)^5^. **a** Single AA substitutions, single AA insertions and N-terminal multi-AA deletions: color intensity indicates the percentage of NS+ (blue) or NS- (red) mutations at each position or losing each position (for multi-AA deletions) (FDR=0.1). **b** Single AA deletions and N-terminal truncations: color intensity depicts the nucleation score of each single AA deletion or of the N-terminal truncation starting at that position. White depicts positions that are not mutated in each dataset.

The DIM dataset also provides substantial mechanistic insight into amyloid nucleation. The vast majority of mutations of any type in the aliphatic C-terminus of Aß strongly reduce aggregation, which is consistent with this region forming the core of all known mature Aß fibril polymorphs ^5,35–39^. The exquisite sensitivity of this region to mutation suggests very strong structural constraints for amyloid nucleation. Only a few specific substitutions and insertions are tolerated, with these concentrated in residues at the end of the peptide which may more easily accommodate different side chains. In addition, the only internal multi-AA deletions that still nucleate despite removing residues from the C-terminus core are those which replace the missing residues with a similar number of aliphatic AA from a more N-terminal region of the peptide. The 14AA aliphatic core of Aß, however, nucleates very poorly unless a positively charged residue is added at the N-terminus. We speculate that this charge may help solubilise this very hydrophobic peptide, prevent an alternative off-pathway non-amyloid aggregation process or participate directly in the nucleation transition state, for example by the formation of a salt bridge with the carboxy-terminus^44^.

In contrast, mutations in the polar N-terminus have much more diverse effects, with 267 variants in this region accelerating aggregation. Much of this region remains unstructured in mature Aß fibrils, including those in AD amyloid plaques (Fig. 5 and Supplementary Fig. 14)^5,35–39^. Mutational effects in this region are not predicted by existing computational methods and they are not obviously interpretable using current mechanistic models of amyloid nucleation: polar, small and positively charged residues as well as P tend to increase nucleation whereas hydrophobic and negatively charged residues tend to decrease it. However, these preferences can also be quite different at individual sites and in sub-regions of the N-terminus, further highlighting the importance of generating complete datasets for the interpretation of clinical variants.

We speculate that mutations in the polar N-terminus of Aß may control the rate of nucleation either because of effects on the ensemble of soluble Aß or because the region participates in the transition state of the nucleation reaction in an as yet undefined manner. Future work should aim to distinguish between these possibilities. The very strong enrichment of disease-causing and aggregation-promoting mutations in the polar N-terminus of Aß makes understanding how polar extensions promote and prevent amyloid aggregation one of the highest priority goals in AD and amyloid research.

## Acknowledgments

M.S. is supported by a fellowship from Agencia de Gestio d’Ajuts Universitaris i de Recerca (2019FI_B 01311). Work in the lab of BB and BL is supported by the la Caixa Research Foundation project ‘DeepAmyloids’ (LCF/PR/HR21/52410004). Work in the lab of BB is also supported by the Spanish Ministry of Science, Innovation and Universities (RTI2018-101491-A-I00 (MICIU/FEDER)) and the CERCA Program/Generalitat de Catalunya. Work in the lab of BL is also supported by a European Research Council (ERC) Advanced Grant (‘Mutanomics’ 883742), the Spanish Ministry of Science, Innovation and Universities (PID2020-118723GB-I00), the Bettencourt Schueller Foundation, the AXA Research Foundation, Agencia de Gestio d’Ajuts Universitaris i de Recerca (AGAUR, 2017 SGR 1322) and the CERCA Program/Generalitat de Catalunya. We also acknowledge the support of the Spanish Ministry of Science and Innovation to the EMBL partnership and the Centro de Excelencia Severo Ochoa. We thank the Chernoff lab for providing strains and plasmids and the CRG Genomics Core Technology for sequencing. We also thank Andre Faure and Marta Badia for advice on data analysis, Leire Moriones for assistance with validation experiments and Xavier Salvatella for discussion.

## Author contributions

M.S. performed all experiments and analyses. M.S., B.L. and B.B. designed the experiments and analyses and wrote the manuscript.

## Supplementary figures and tables

**Supplementary Figure 1. Reproducibility and assay validation**

**a** Correlation of nucleation scores for three biological replicates (n_1-2_=2,951, n_1-3_=2,984, n_2-3_=2,950 genotypes). **b** Correlation of nucleation scores measured for the synthetic library used in this study and a previous library generated by error-prone PCR (n=423 common variants)^15^. **c** Correlation of nucleation scores measured in the competition experiment or individually for selected variants (n=10). Vertical and horizontal error bars indicate 95% confidence intervals of mean NS. Pearson correlation coefficients are indicated in (**a-c**). **d** Correlation of nucleation scores with *in vitro* primary and secondary nucleation rate constants^45^. Weighted Pearson correlation coefficients are indicated. **e** Receiver operating characteristic (ROC) curves for 12 of all the single AA substitutions described as dominant fAD variants (H6R, D7N, D7H, E11K, K16Q, A21G, E22Q, E22K, E22G, D23N, L34V and A42T) versus all other single AA substitutions present in the dataset (n_non-fAD_=739) for two DMS datasets (Nucleation score and Solubility score), aggregation predictors (Tango, Zyggregator, Waltz, Camsol^29–32^) and variant effect predictors (Polyphen and CADD^46,47^). Area under the curve (AUC) values are indicated. Diagonal dashed line indicates the performance of a random classifier. **f** Number and type of variants increasing nucleation (NS-, FDR=0.1) for each peptide region.

**Supplementary Figure 2. Mutational effects of single AA substitutions and insertions**

**a** Heatmap of nucleation scores FDR categories for single AA substitutions. The WT AA and position are indicated in the x-axis and the mutant AA is indicated in the y-axis. Variants not present are represented in gray. Synonymous mutants are indicated with ‘*’ and fAD mutants with a black box. **b** Heatmap of nucleation scores FDR categories for single AA insertions. **c** Frequency of increasing or decreasing nucleation (FDR=0.1) single AA substitutions upon substituting specific WT AA, for each peptide region.

**Supplementary Figure 3. Mutational effects of single AA substitutions and insertions**

**a,b** Clustering of single AA mutation nucleation scores by mutated residue identity and position. Position is indicated in the x-axis; AA insertions were considered after (**a**) or before (**b**) each position. Mutations are indicated in the y-axis and labeled with an ‘s’ for substitutions or an ‘i’ for insertions, followed by the substituted or inserted AA. ‘del’ indicates single AA deletion of that position.

**Supplementary Figure 4. Comparing the mutational effects of single AA variants**

**a** Correlation of average nucleation scores for each position, for single AA insertions before or after a specific position and single AA substitutions (left) or single AA deletions (middle), and for single AA deletions and single AA substitutions (right) at the corresponding position. Color code indicates peptide region (N-terminus, AA 1-28, or C-terminus, AA 29-42). **b** Correlation of average nucleation scores for each AA, for single AA deletions and single AA substitutions (top row), single AA insertions and single AA substitutions (middle row) and single AA insertions and single AA deletions (bottom row); and for the full peptide (left column), the N-terminus (AA 1-28, middle column) or the C-terminus (AA 29-42, right column). **c** Correlation of average nucleation scores for each AA, for the C and the N-terminus, for single AA substitutions (left), single AA insertions (middle) and single AA deletions (right). AA labels are coloured by AA class in (**b**) and (**c**). Pearson correlation coefficients are indicated. Dashed lines indicate the WT nucleation score (0).

**Supplementary Figure 5. Comparing the mutational effects of single AA substitutions and insertions**

**a,b** Correlation of nucleation scores at each position arranged by each AA type, between single AA substitutions and single AA insertions before (**a**) or after (**b**) the corresponding position. Pearson correlation coefficients are indicated. Dashed lines indicate the WT nucleation score (0).

**Supplementary Figure 6. Comparing the mutational effects of single AA substitutions and insertions**

**a,b** Correlation of nucleation scores for each AA type arranged by position, between single AA substitutions and single AA insertions before (**a**) or after (**b**) the corresponding position. Pearson correlation coefficients are indicated. Dashed lines indicate the WT nucleation score (0). Color code indicates AA position.

**Supplementary Figure 7. Mutational effects of substituting in specific AAs**

The wild-type (WT) AA and position are indicated on the x-axis and coloured on the basis of their effect (NS+ or NS-) and FDR category. The horizontal line indicates the WT nucleation score (0).

**Supplementary Figure 8. Mutational effects of inserting specific AAs**

The wild-type (WT) AA and position are indicated on the x-axis and coloured on the basis of their effect (NS+ or NS-) and FDR category. The horizontal line indicates the WT nucleation score (0).

**Supplementary Figure 9. Evaluation of mutational effect and aggregation predictors**

**a** Correlation of nucleation scores with the predictions of aggregation predictors (Tango, Zyggregator, Waltz and Camsol)^29–32^, variant effect predictors (CADD, Polyphen)^46,47^, solubility scores^48^, PC1^49^ and hydrophobicity^50^ for single AA mutations, at the N-terminus (left) or the C-terminus (right). Pearson correlation coefficients are indicated. Dashed lines indicate the WT nucleation score (0). **b** Receiver operating characteristic (ROC) curves for classifying increasing nucleation variants (NS+, FDR=0.1) for single AA mutations, at the N and C-terminus, for aggregation predictors^29–32^, variant effect predictors^46,47^, solubility scores^48^, PC1^49^ and hydrophobicity^50^. Area under the curve (AUC) values are indicated. Diagonal dashed line indicates the performance of a random classifier.

**Supplementary Figure 10. Multi-AA deletions**

**a** Heatmap of nucleation scores FDR categories for multi-AA deletions. The WT AA and position of the first and last residues deleted are indicated in the x-axis and y-axis, respectively. The black squares indicate fAD variants: Osaka (E22Δ) and Uppsala (Δ19-24). Variants not present are represented in gray. **b** Effect on nucleation of variants that delete E22, D23N or both. The distance the closest negative residue (D,E) - if present - to the C-terminus (AA 29-42) is shown in the x-axis. Variants with no negative residues are also shown (no D/E). Shape indicates the identity of the residue and color code indicates FDR=0.1 category. The horizontal line indicates the WT nucleation score (0). **c** Generation of new N-terminus sequences flanking the Aß core. Nucleation score distributions of each AA at each position for deletions at the N-terminus. Distance from the C-terminus (AA 29-42) is indicated in the x-axis, as well as WT AA and position. Color of the violin plot indicates median nucleation score for each distribution.

**Supplementary Figure 11. Multi-AA deletion variants**

**a** AA sequence for variants with internal multi-AA deletions located at both N and C-terminus, with significantly decreased nucleation (FDR=0.1). **b** AA sequence for deletions at the N-terminus removing residue K28, with positive nucleation score (NS>0).

**Supplementary Figure 12. N- and C-terminal truncations**

**a,b** Heatmap of nucleation scores (**a**) or FDR categories (**b**) for truncations from one or both ends of the peptide. The WT AA and position of the first and last residues of the resulting peptide are indicated in the x-axis and y-axis, respectively.

**Supplementary Figure 13. Top nucleating sequences in the library**

AA sequence for 1% variants with highest NS (all FDR=0.1) in the library. AA are coloured by AA class.

**Supplementary Figure 14**. **Impact of diverse classes of mutations along the structure of Aß42 fibrils from sporadic AD brains**

The impact of all mutations of all classes is summarized over the structure of Aß42 fibrils (PDB: 7Q4B)^5^. In fibrils extracted from sporadic AD brains, Aß42 adopts a S-shaped structure at the C-terminus with an N-terminal arm linking to an unstructured region. **a** Single AA substitutions, single AA insertions and N-terminal multi-AA deletions: Color intensity indicates the percentage of NS+ (blue) or NS- (red) mutations at each position or losing each position (for multi-AA deletions) (FDR=0.1). **b** Single AA deletions and N-terminal truncations: Color intensity depicts the nucleation score of each single AA deletion or of the N-terminal truncation starting at that position. White depicts positions that are not mutated in each dataset.

**Supplementary Table 1**

List of candidate fAD pathogenic variants with increased nucleation (FDR=0.1).

**Supplementary Table 2**

List of N-terminal Aß truncations reported in the literature and their corresponding nucleation score and category.

**Supplementary Table 3**

List of oligonucleotides used in this study.

**Supplementary Table 4**

Processed data required to reproduce the analysis and figures in this paper, with read counts, nucleation scores, FDR category, associated error terms and associated pathogenicity.

## Material and Methods

### Library design

The designed library contains a total of 3,164 unique Aß42 variants, with all single AA substitutions at each position (n=798), all single AA insertions at all positions (n=780), all deletions ranging from 1 to 39 AA in size in all positions (n=768), sequences truncated from either one or both ends of the peptide with a minimum peptide length of 3 AA and maximum peptide length of 40 AA (n=817), and the Aß42 WT sequence (n=1).

### Plasmid Library construction

The synthetic library was synthesized by Twist Bioscience and consisted of an Aß42 variant region of 9 nt to 129 nt, flanked by 25 nt upstream and 21 nt downstream constant regions. 10ng of the library were amplified by PCR (Q5 high-fidelity DNA polymerase, NEB) for 12 cycles with primers annealing to the constant regions (primers MS_01-02, Supplementary Table 3), according to the manufacturer’s protocol. The product was then purified by column purification (MinElute PCR Purification Kit, Qiagen). In parallel, the P_CUP1_-Sup35N-Aß42 plasmid was linearised by PCR (Q5 high-fidelity DNA polymerase, NEB) with primers that remove the WT Aß42 sequence (primers MS_03-04, Supplementary Table 3). The product was purified from a 1% agarose gel (QIAquick Gel Extraction Kit, Qiagen).

The library was then ligated into 100ng of the linearised plasmid in a 5:1 (insert:vector) ratio by a Gibson approach with 3h of incubation followed by dialysis for 45 min on a membrane filter (MF-Millipore 0,025 um membrane, Merck). The product was transformed into 10-beta Electrocompetent E.coli (NEB), by electroporation with 2.0kV, 200 Ω, 25 uF (BioRad GenePulser machine). Cells were recovered in SOC medium for 30 min and grown overnight in 30 ml of LB ampicillin medium. A small amount of cells were also plated in LB ampicillin plates to assess transformation efficiency. A total of 50,000 transformants were estimated, meaning that each variant in the library is represented >15 times. 5ml of overnight culture were harvested to purify the Aß42 library with a mini prep (QIAprep Miniprep Kit, Qiagen).

### Yeast transformation

*Saccharomyces cerevisiae* [psi-pin-] (MATa ade1-14 his3 leu2-3,112 lys2 trp1 ura3-52) strain was used in all experiments in this study.

Yeast cells were transformed with the Aß42 plasmid library in three biological replicates. An individual colony was grown overnight in 25ml YPDA medium at 30 C and 200 rpm. Cells were diluted in 150 ml to OD600 =0.25 and grown for 4-5 h. When cells reached the exponential phase (OD~0.7-0.8), they were splitted in 10 transformation tubes of 15 ml each. Each tube was treated as follows: cells were harvested at 400 g for 5 min, washed with milliQ and resuspended in 1 ml YTB (100 mM LiOAc, 10 mM Tris pH 8.0, 1 mM EDTA). They were harvested again and resuspended in 72 ul YTB. 100 ng of plasmid library were added to the cells, together with 8 ul of salmon sperm DNA (UltraPure, Thermo Scientific) previously boiled, 60 ul of dimethyl sulfoxide (Merck) and 500 ul of YTB-PEG (100 mM LiOAc, 10 mM Tris pH 8.0, 1 mM EDTA, 40% PEG 3350). Heat-shock was performed at 42 C for 14 min in a thermo block. Finally, cells were harvested and resuspended in 300 ml plasmid selection medium (-URA, 20% glucose), pooling together the 10 transformation tubes and allowing them to grow for 50 h at 30 C and 200 rpm. A small amount of cells were also plated in plasmid selection medium to assess transformation efficiency. A total of 118,125, 152,000 and 139,500 transformants were estimated for each biological replicate respectively, meaning that each variant in the library is represented >37 times.

After 50 h, cells were diluted in 25 ml plasmid selection medium to OD =0.02 and grown exponentially for 15 h. Finally, the culture was harvested and stored at −80 C in 25 % glycerol.

### Selection experiments

*In vivo* selection assays were performed in five technical replicates for each biological replicate. For each technical replicate, cells were thawed from −80 C in 20 ml plasmid selection medium at OD=0.05 and grown until exponential for 15 h. At this stage, cells were harvested and resuspended in 20 ml protein induction medium (-URA, 20% glucose, 100 uM Cu2SO4) at OD=0.05. After 24 h the 4x 5ml input pellets were collected and 1 million cells/replicate were plated on-ADE-URA selection medium in 145 cm^2^ plates (Nunc, Thermo Scientific). Plates were incubated at 30 C for 7 days inside an incubator. Finally colonies were scraped off the plates with PBS 1x and harvested by centrifugation to collect the output pellets. Both input and output pellets were stored at −20 C for later DNA extraction.

### DNA extraction and sequencing library preparation

One input and one output pellets for each technical and biological replicate (2×5×3 samples) were resuspended in 0.5 ml extraction buffer (2% Triton-X, 1% SDS, 100mM NaCl, 10mM Tris-HCl pH8, 1mM EDTA pH8). They were then freezed for 10 min in an ethanol-dry ice bath and heated for 10 min at 62 C. This cycle was repeated twice. 0.5 ml of phenol:chloroform:isoamyl (25:24:1 mixture, Thermo Scientific) was added together with glass beads (Sigma). Samples were vortexed for 10 min and centrifuged for 30 min at 4000 rpm. The aqueous phase was then transferred to a new tube, and mixed again with phenol:chloroform:isoamyl, vortexed and centrifuged for 45 min at 4000 rpm. Next, the aqueous phase was transferred to another tube with 1:10V 3M NaOAc and 2.2V cold ethanol 96% for DNA precipitation. After 30min at −20 C, samples were centrifuged and pellets were dried overnight. The following day, pellets were resuspended in 0.3 ml TE 1X buffer and treated with 10ul RNAse A (Thermo Scientific) for 30 min at 37 C. DNA was finally purified using 10 ul of silica beads (QIAEX II Gel Extraction Kit, Qiagen) and eluted in 30 ul elution buffer. Plasmid concentrations were measured by quantitative PCR with SYBR green (Merck) and primers annealing to the origin of replication site of the P_CUP1_-Sup35N-Aß42 plasmid at 58 C for 40 cycles (primers MS_05-06, Supplementary Table 3).

The library for high-throughput sequencing was prepared in a two-step PCR (Q5 high-fidelity DNA polymerase, NEB). In PCR1, 50 million of molecules were amplified for 15 cycles with frame-shifted primers with homology to Illumina sequencing primers (primers MS_07-20, Supplementary Table 3). The products were purified with ExoSAP treatment (Affymetrix) and by column purification (MinElute PCR Purification Kit, Qiagen). They were then amplified for 10 cycles in PCR2 with Illumina indexed primers (primers MS_21-37, Supplementary Table 3). The six samples of each technical replicate were pooled together equimolarly and the final product was purified from a 2% agarose gel with 20 ul silica beads (QIAEX II Gel Extraction Kit, Qiagen).

The library was sent for 125 bp paired-end sequencing in an Illumina HiSeq2500 sequencer at the CRG Genomics core facility. In total, >426 million paired-end reads were obtained, which is between 7-20 million per sample (i.e. input or output for a specific technical and biological replicate), representing >2200x read coverage for each designed variant in the library.

### Individual variant testing

Selected Aß42 variants for individual testing were obtained by PCR linearisation (Q5 high-fidelity DNA polymerase, NEB) with mutagenic primers (primers MS_38-47, Supplementary Table 3). PCR products were treated with Dpn1 overnight and transformed in DH5α competent E.coli. Plasmids were purified by mini prep (QIAprep Miniprep Kit, Qiagen) and transformed into yeast cells using one transformation tube of the transformation protocol described above. All constructions were verified by Sanger sequencing.

Yeast cells expressing individual variants were grown overnight in plasmid selection medium (-URA 20% glucose). They were then diluted to OD 0.05 in protein induction medium (-URA 20% glucose 100uM Cu2SO4) and grown for 24h. Cells were plated on -URA (control) and -ADE-URA (selection) plates in three independent replicates, and allowed to grow for 7 days at 30 C. Adenine growth was calculated as the percentage of colonies in -ADE-URA relative to colonies in -URA.

### Data processing

FastQ files from paired end sequencing of the Aß42 library were processed using the DiMSum pipeline (https://github.com/lehner-lab/DiMSum)25, an R package that wraps sequencing processing tools, such as FastQC (http://www.bioinformatics.babraham.ac.uk/projects/fastqc/) for quality assessment; Cutadapt^51^ for constant region trimming; and VSEARCH^52^ for read alignment. 5’ and 3’ constant regions were trimmed, allowing a maximum of 20% of mismatches relative to the reference sequence. Sequences with a Phred base quality score below 30 were discarded. At this stage, around 370 million reads passed the filtering criteria.

Unique variants were then aggregated and counted using Starcode (https://github.com/gui11aume/starcode). Non-designed variants were also discarded for further analysis, as well as variants with less than 10 input reads in any of the replicates and variants resulting from one single nt change with less than 1000 input reads. Estimates from DiMSum^25^ were used to choose the filtering thresholds.

### Nucleation scores and error estimates

The DiMSum package (https://github.com/lehner-lab/DiMSum)25 was also used to calculate nucleation scores (NS) and their error estimates for each variant in each biological replicate as:

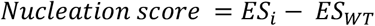

Where *ES_i_* = *log*(*F_i_ OUTPUT*) -*log*(*F_i_ INPUT*) for a specific variant and

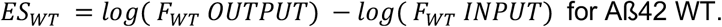

NSs for each variant were merged across biological replicates using error-weighted mean and centered to the WT Aß42 NS. All NS and associated error estimates are available in Supplementary Table 4.

### Data analysis

#### Variants in the library

NS was obtained for 3,087 unique Aß42 variants, which were splitted into mutation classes: 751 single AA substitutions, 763 single AA insertions, 37 single AA deletions, 729 internal multi-AA deletions, 817 truncations (from one or both ends) and WT Aß42.

In addition, nine variants (2 single AA substitutions, 6 single AA insertions and one multi-AA deletion) were classified as non-nucleating but do not have an associated NS (ie. they have input reads but no output reads) and are indicated as such in Figs. 2a,b and Fig. 3a. Each variant is assigned to one mutation class: deletions from position 1 or 42 are classified as truncations and not deletions, and deletions of positions 1 and 42 are classified as single AA deletion and not as truncations. Multiple mutation classes can be combined for visualization or analysis (e.g. truncations and single deletions are included in the deletions matrix in Fig. 3a).

We assign to single AA insertions the position of the inserted AA (e.g. an insertion between positions 1 and 2 is an insertion at position 2). In the case of insertions between positions 28 and 29 (i.e. between the N and C-terminus), they are insertions at position 29 but considered N-terminal mutations.

Different mutations can result in the same coding sequence (e.g. H13Δ and H14Δ, or DAEDVGSNKGAIIGLMVGGVVIA, which is Δ2-20, Δ3-21 and Δ4-22). This is the case for single AA insertions, single and multi-AA deletions. In general, they are only considered as one coding variant but considered multiple times for visualization or if the analysis is position-specific, in figures: Figs. 2b-f, 3a,b and 5, and Supplementary Figs. 2-6 and 10a.

#### Aggregation and variant effect predictors

For the aggregation predictors (Tango, Zyggregator, Waltz, Camsol^29–31^), individual residue-level scores were summed to obtain a score per single AA mutation sequence. We then calculated the log value for each variant relative to the WT score. For the variant effect predictors (Polyphen and CADD^46,47^), we also calculated the log value for each single AA substitutions variant but in this case values were scaled relative to the lowest predicted score.

We also used an hydrophobicity scale^50^ and a principal component from a previous study (PC1^49^) that relates strongly to changes in hydrophobicity. For each single AA substitution variant, the values of a specific AA property represent the difference between the mutant and the WT scores.

#### ROC analysis

ROC curves and AUC values were built and obtained using the ‘pROC’ R package. The table of fAD mutations was taken from https://www.alzforum.org/mutations/app. The nucleation scores and categories for all fAD variants, as well as the criteria used to consider them as fAD, are reported in Supplementary Table 4.

### Data availability

Raw sequencing data and the processed data table (Supplementary Table 4) are deposited in NCBI’s Gene Expression Omnibus (GEO) as GSE193837. All scripts used for downstream analysis and to reproduce all figures are in https://github.com/BEBlab/DIM-abeta.

