## Supplementary figures 1-14 for "An atlas of amyloid aggregation: the impact of substitutions, insertions, deletions and truncations on amyloid beta fibril nucleation"

### Supplementary Fig. 1

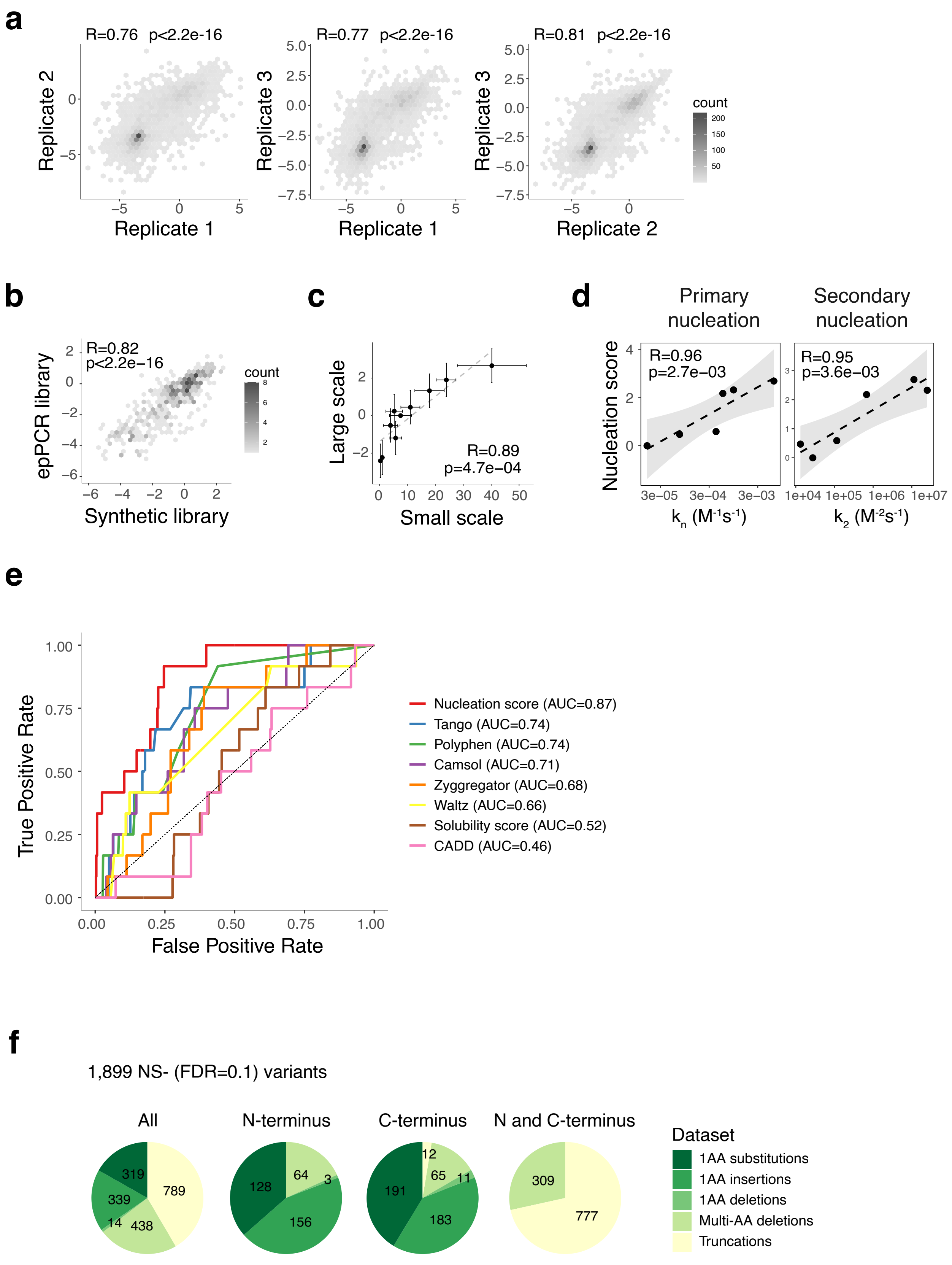

### Supplementary Fig. 2

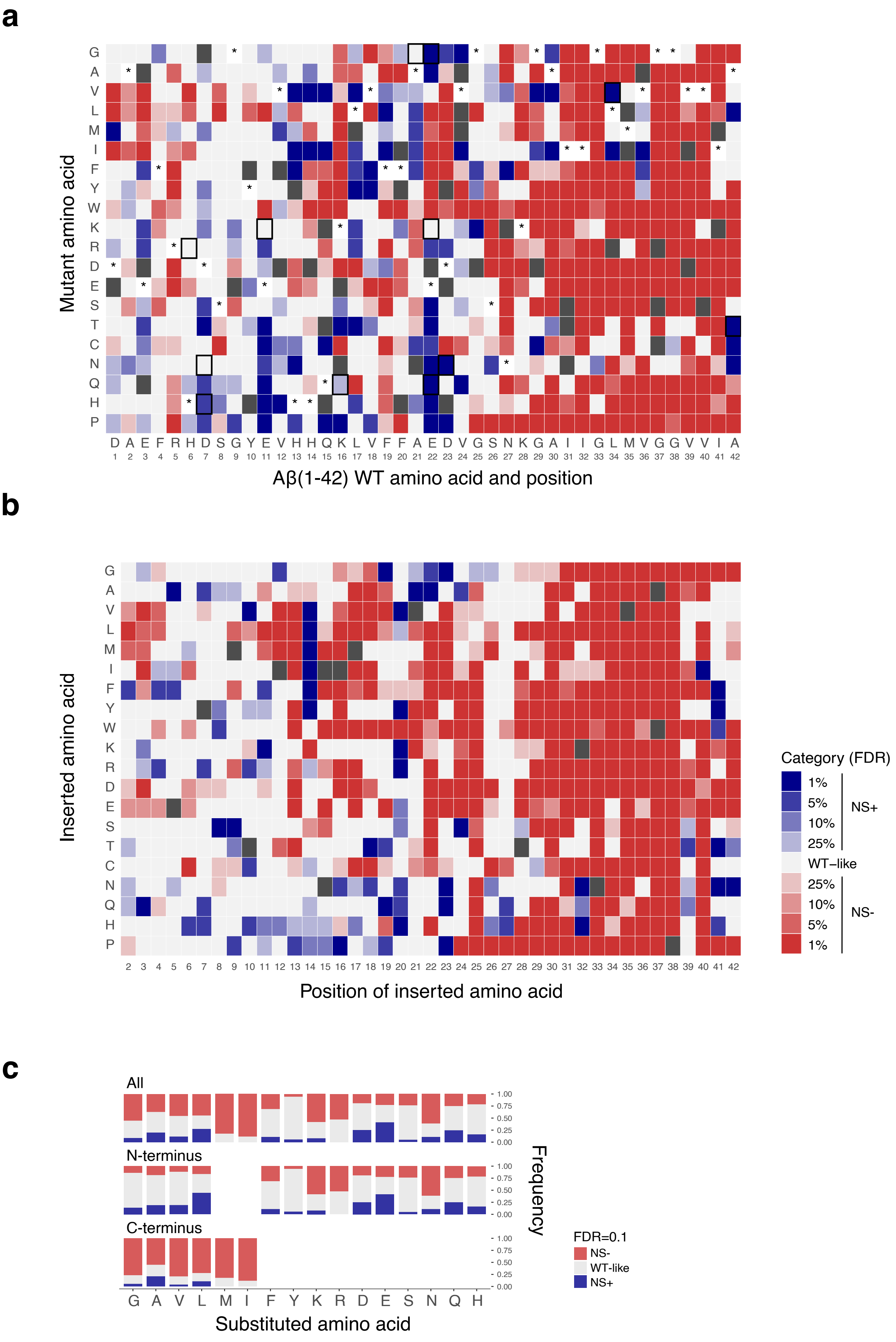

**Supplementary Fig. 3**

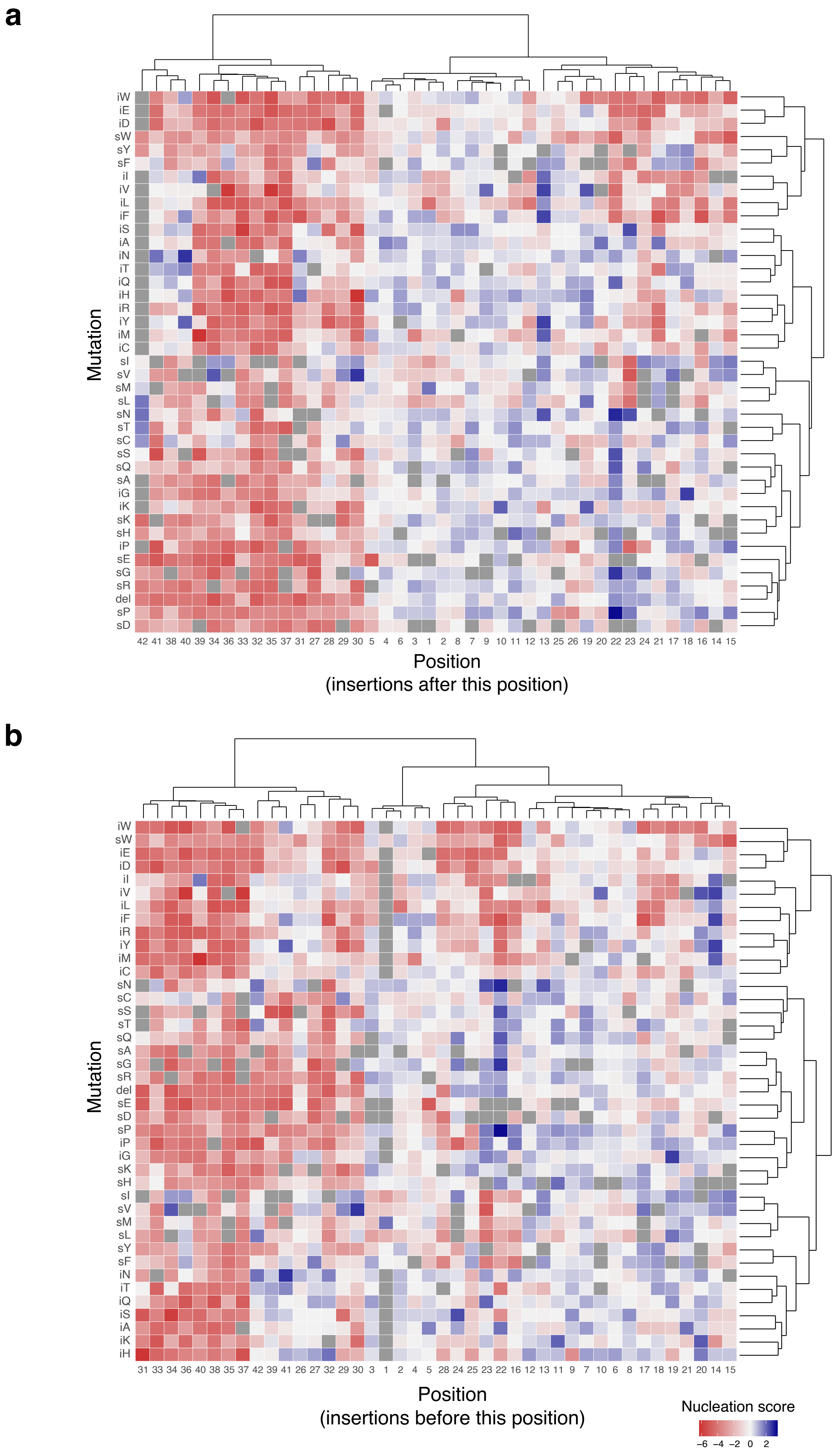

#### Supplementay Fig. 4

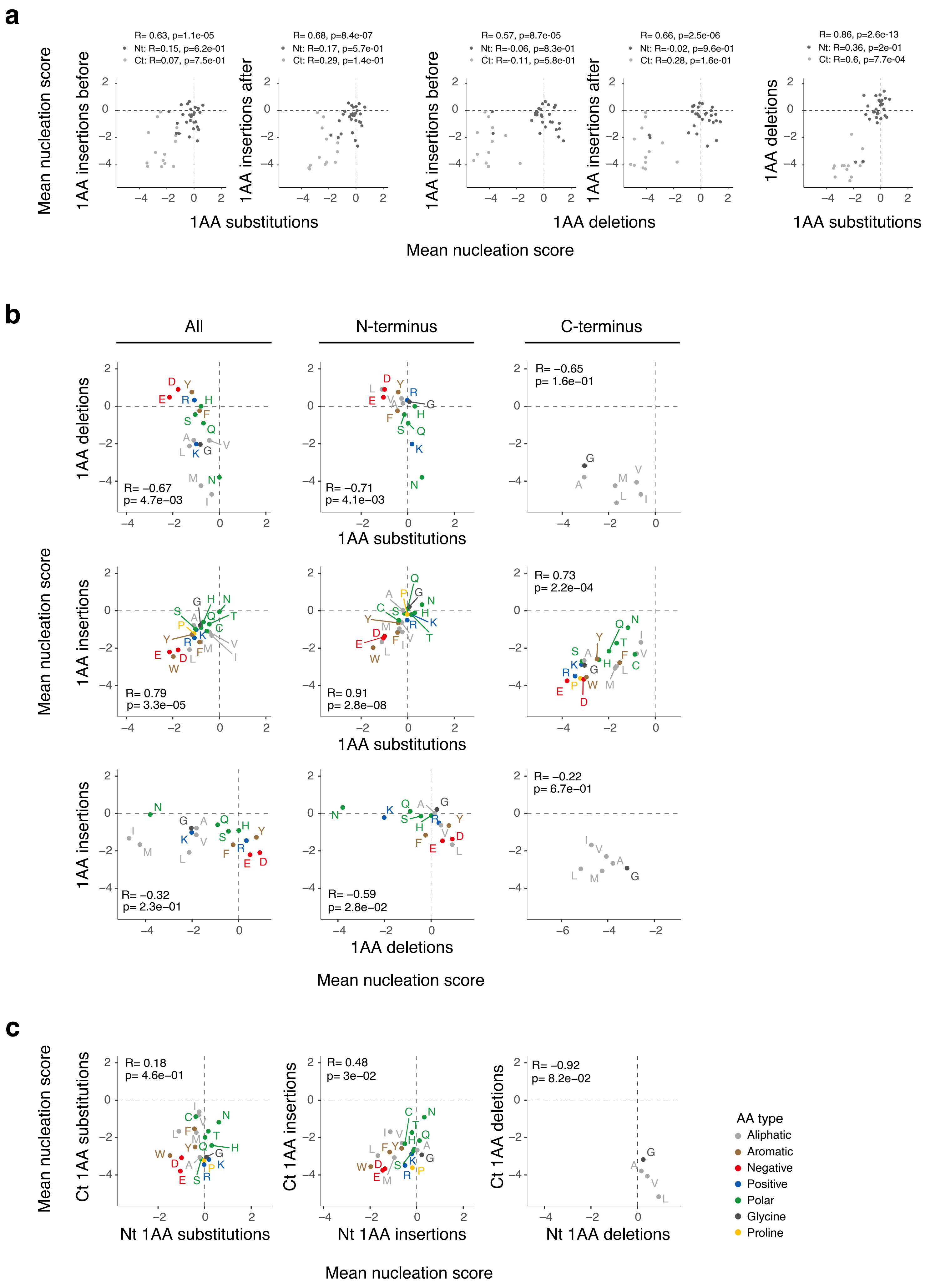

### Supplementary Fig. 5

a

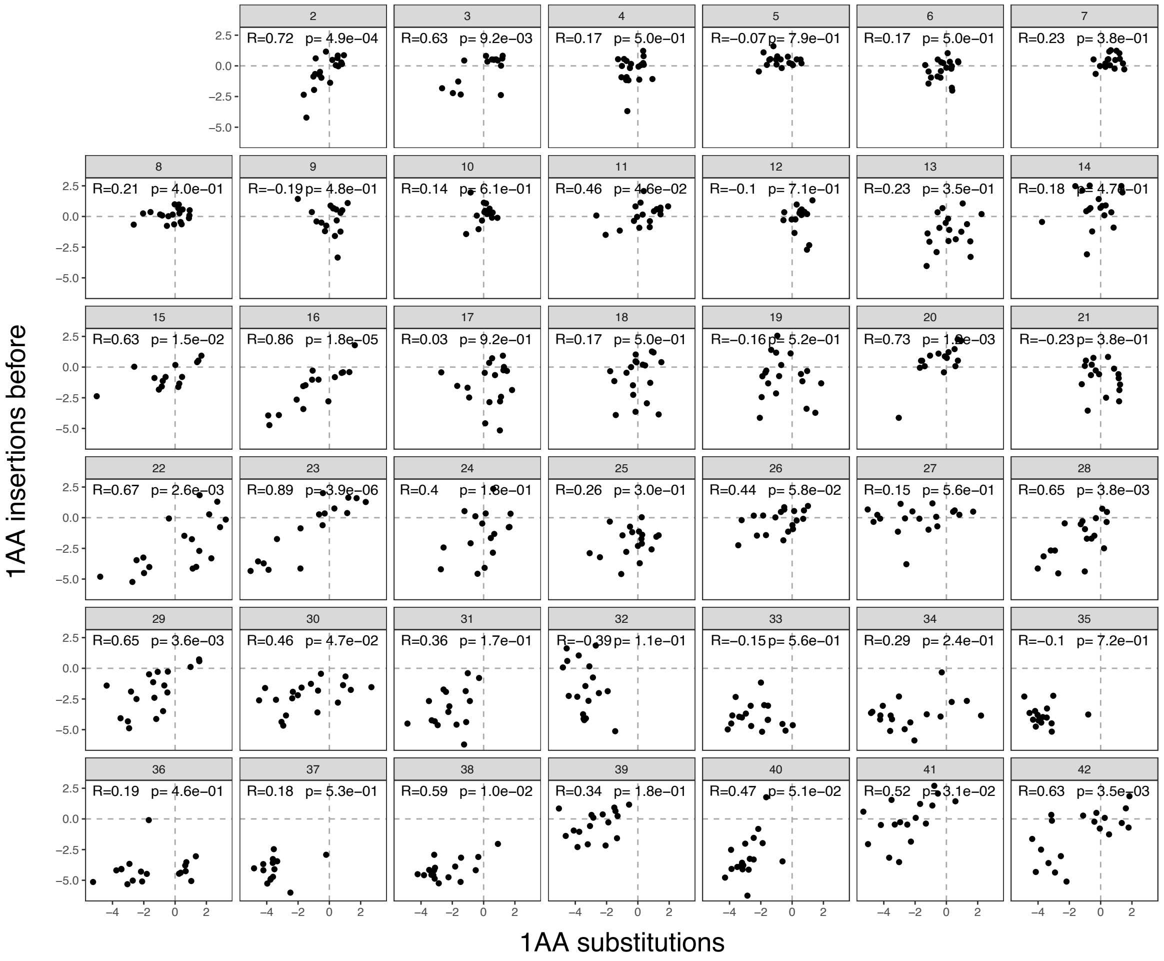


b

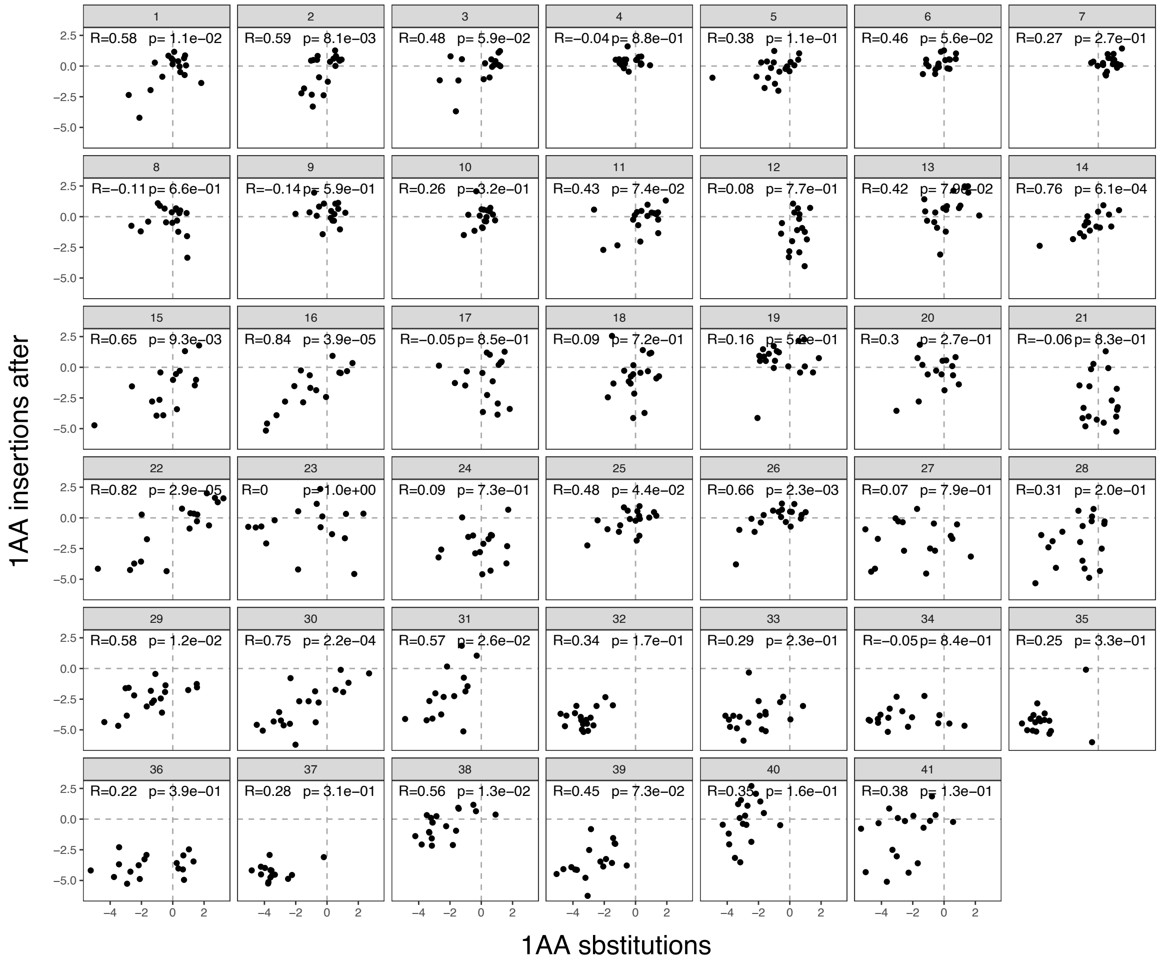

### Supplementary Fig. 6

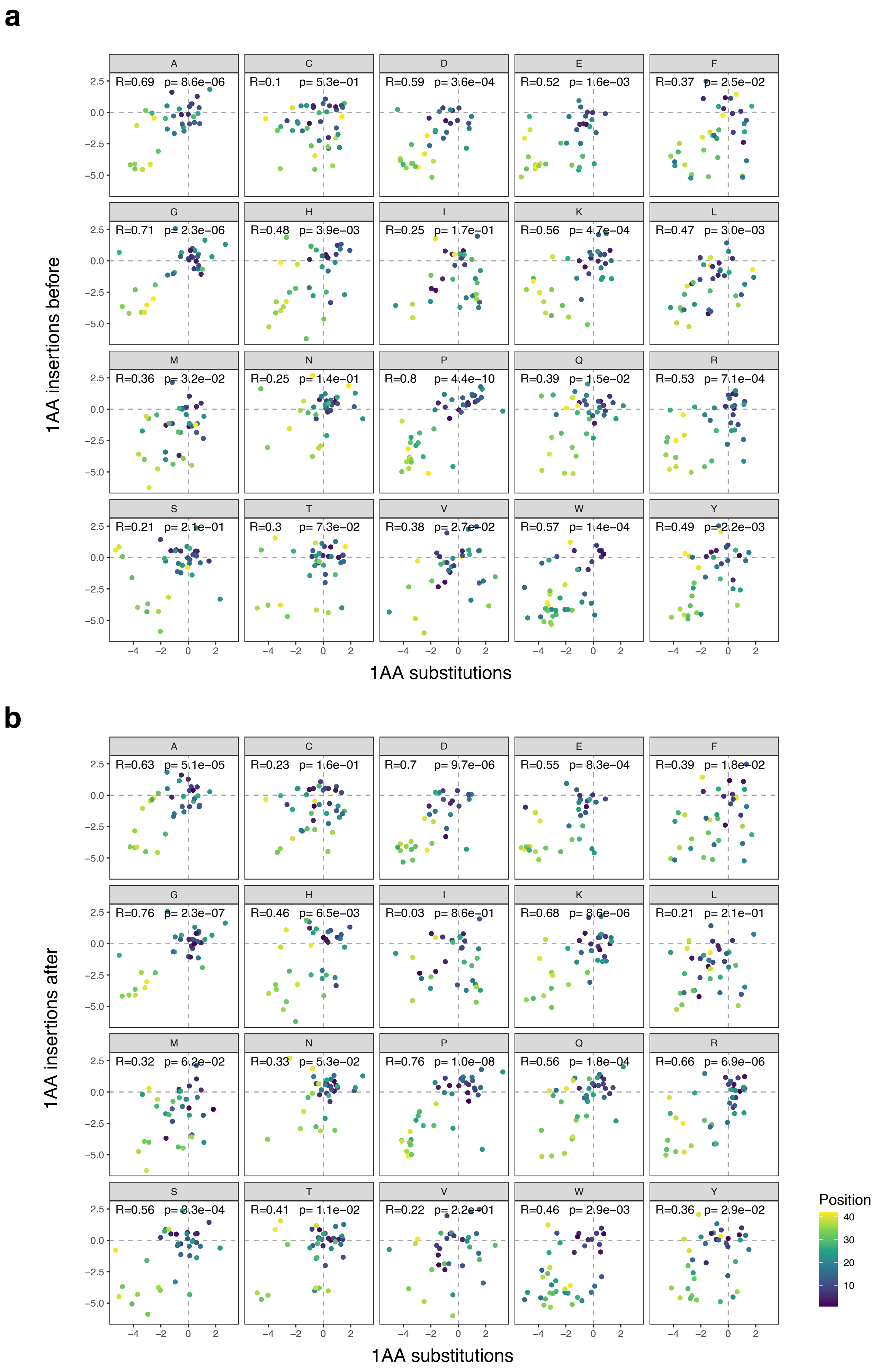

### Supplementary Fig. 7

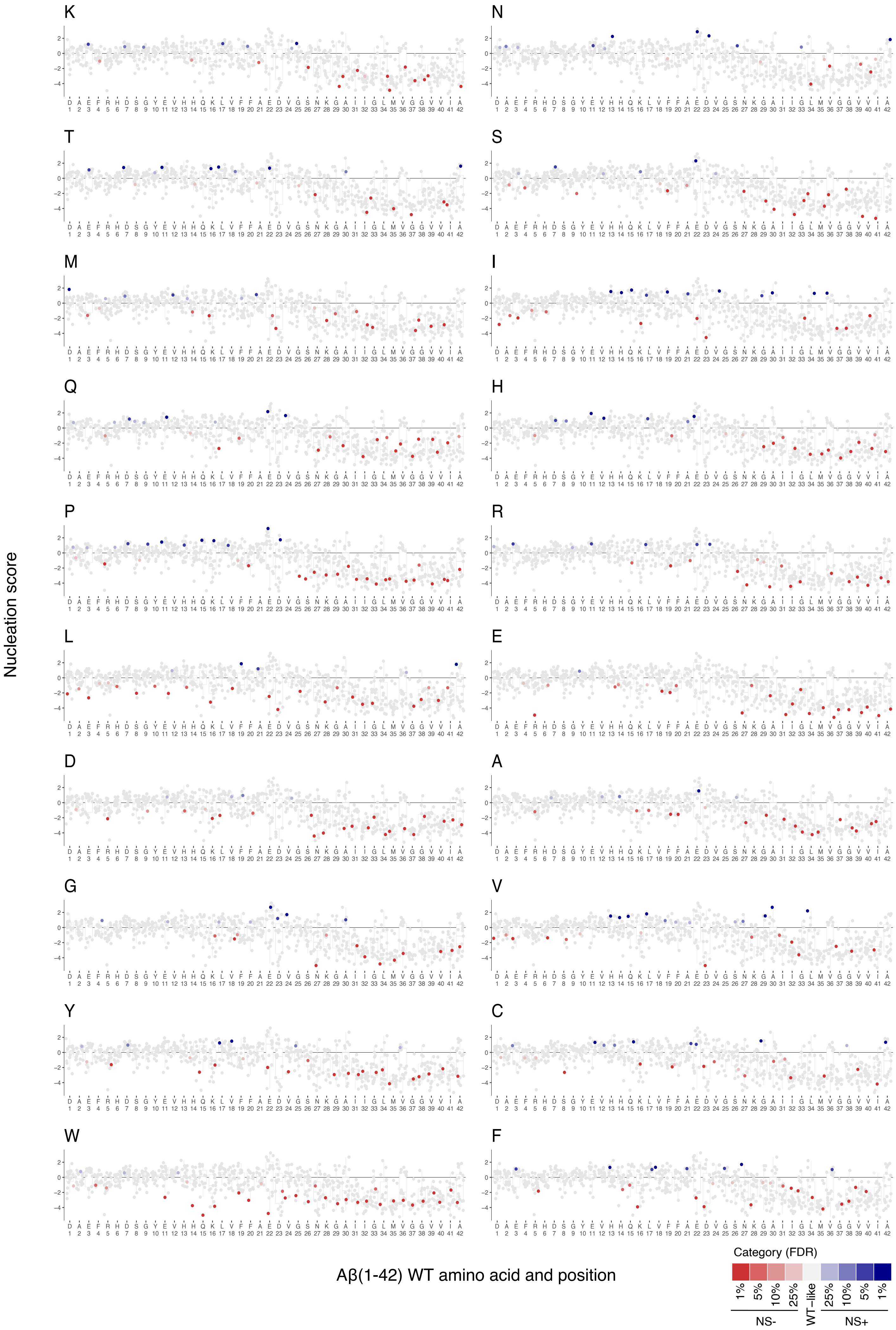

#### Supplementary Fig. 8

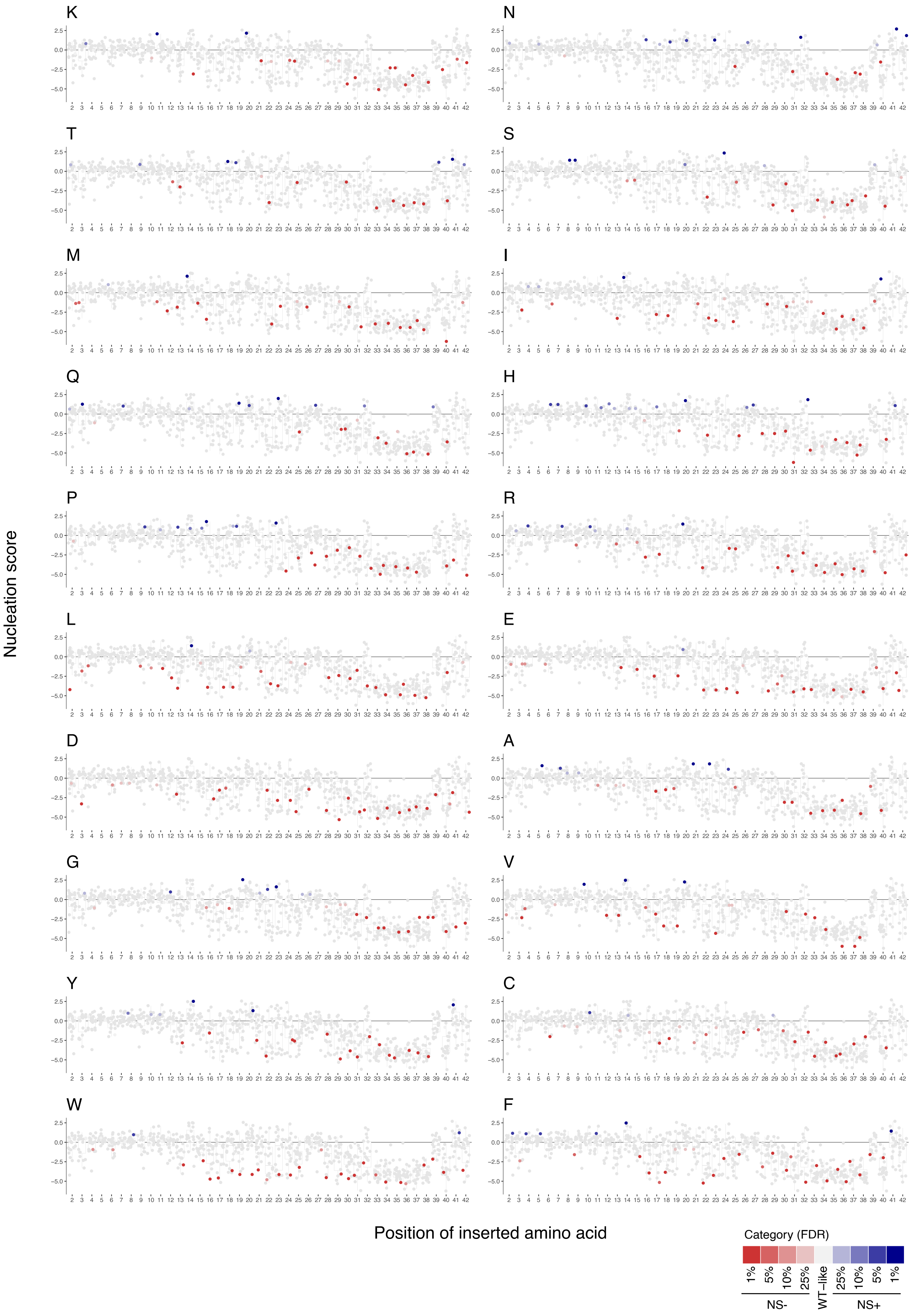

### Supplementary Fig. 9

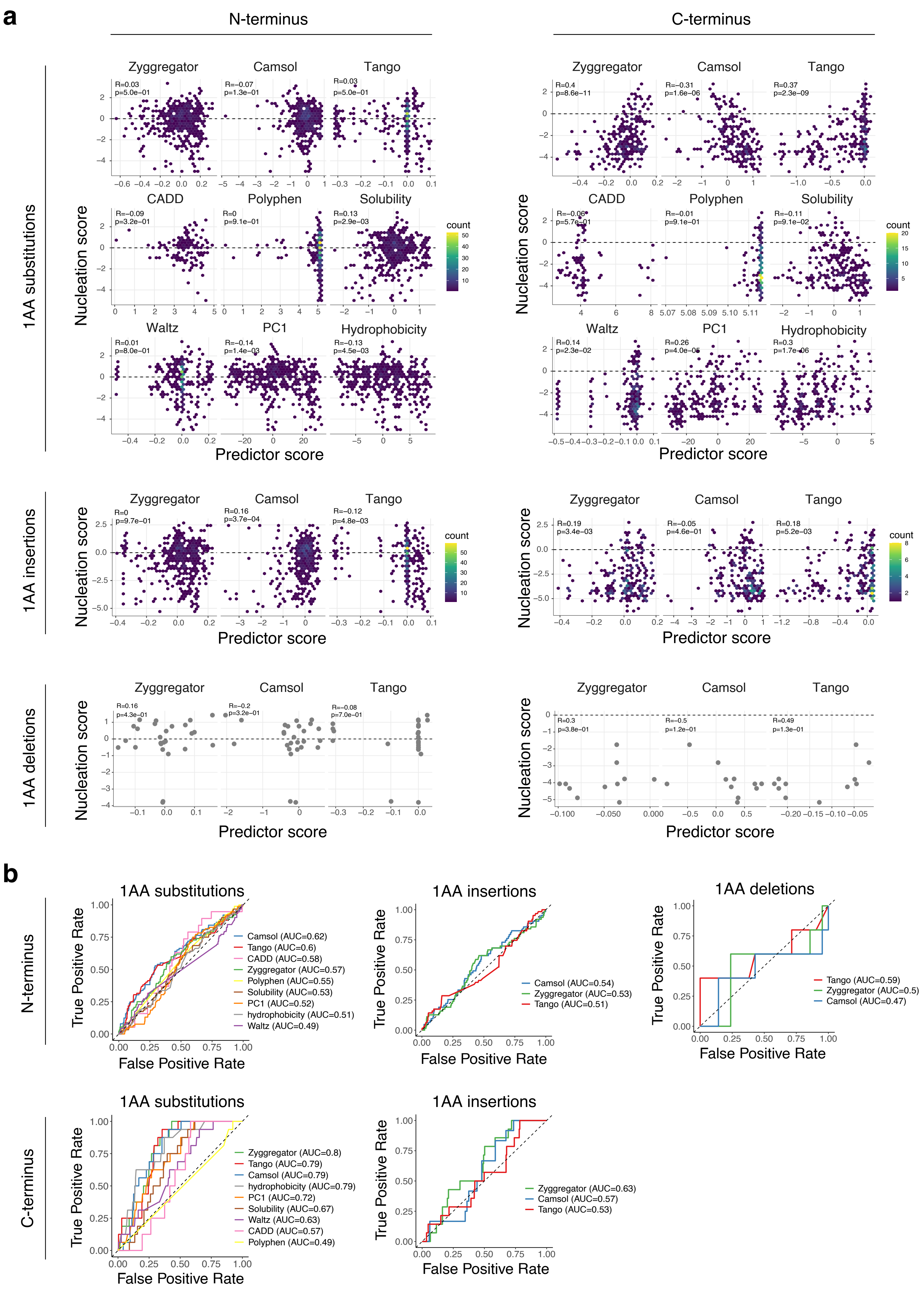

#### Supplementary Fig. 10

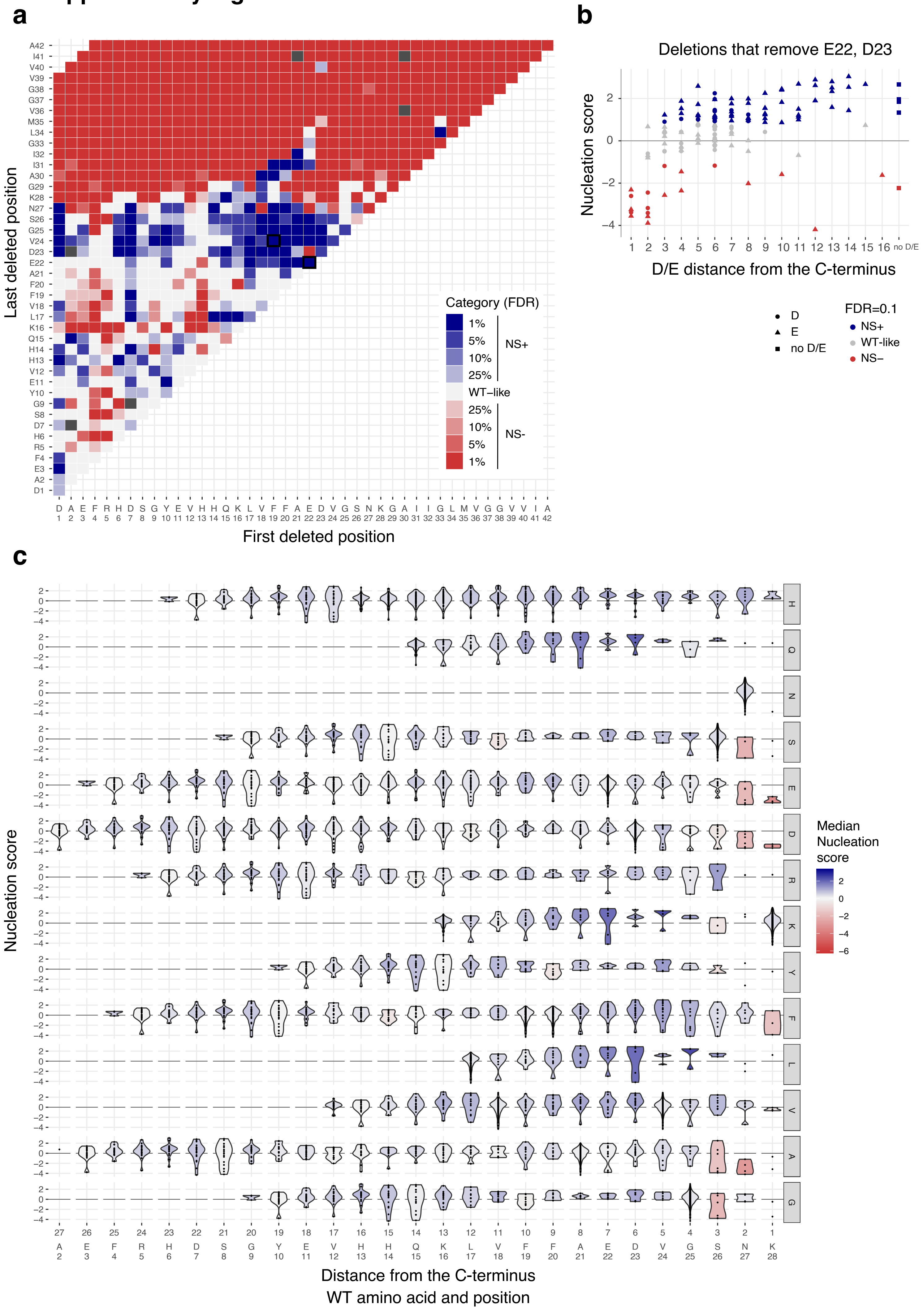

### Supplementary Fig. 11

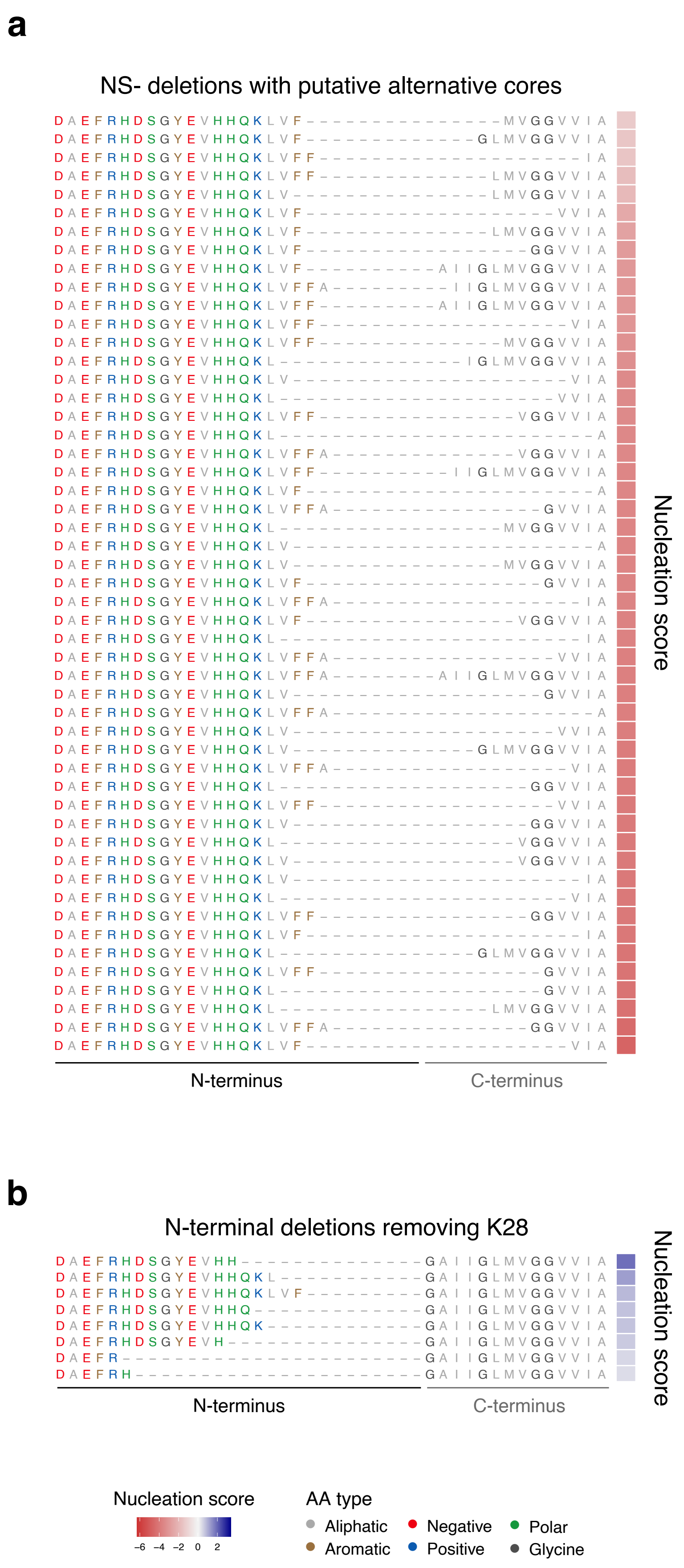

#### Supplementary Fig. 12

**a**

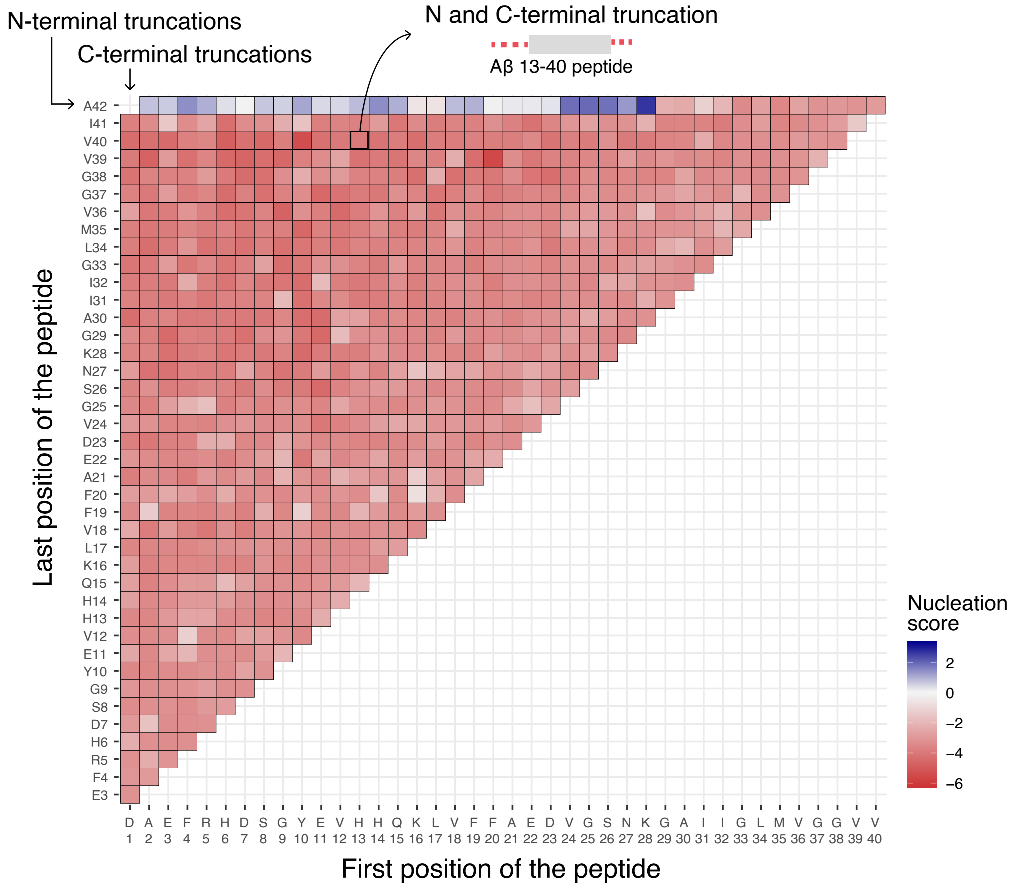


**b**

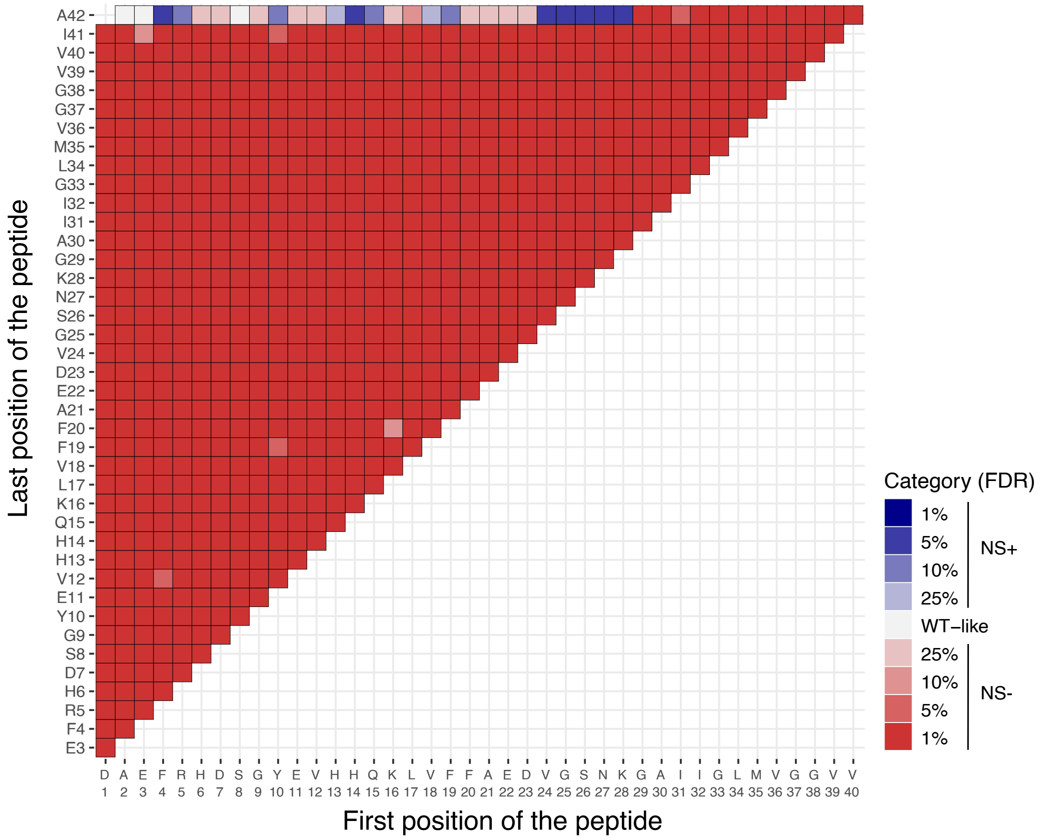

#### Supplementary Fig. 13

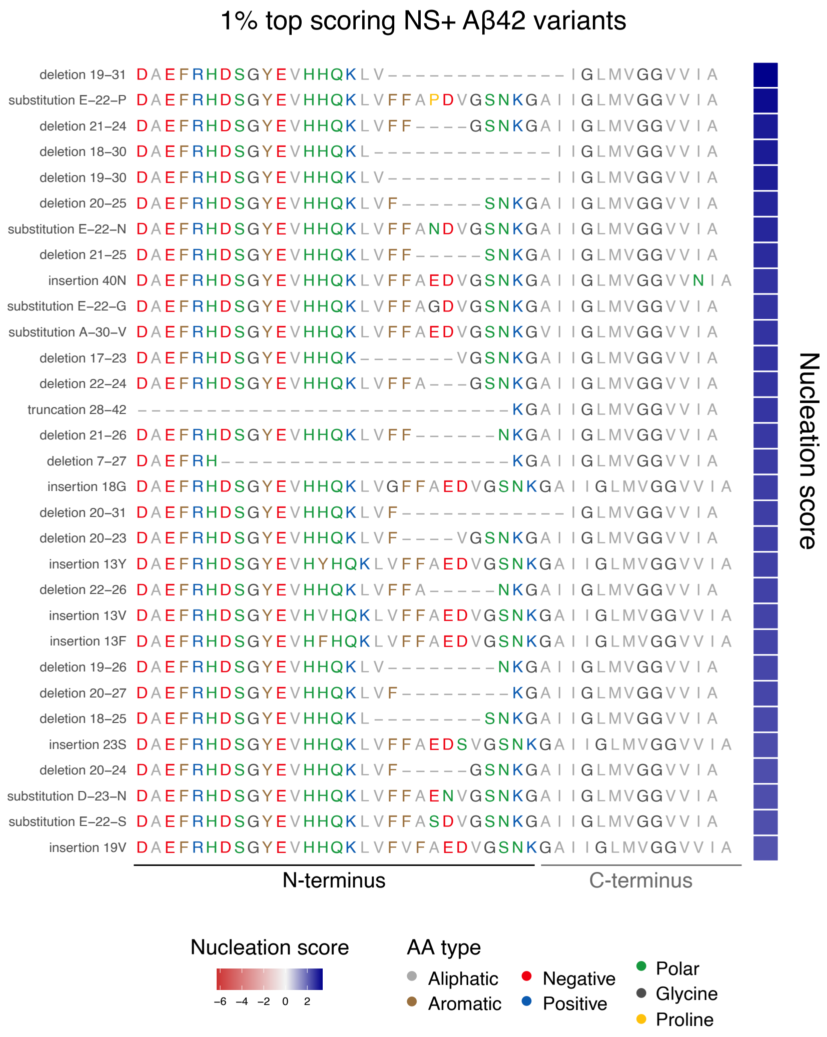

### Supplementary Fig. 14

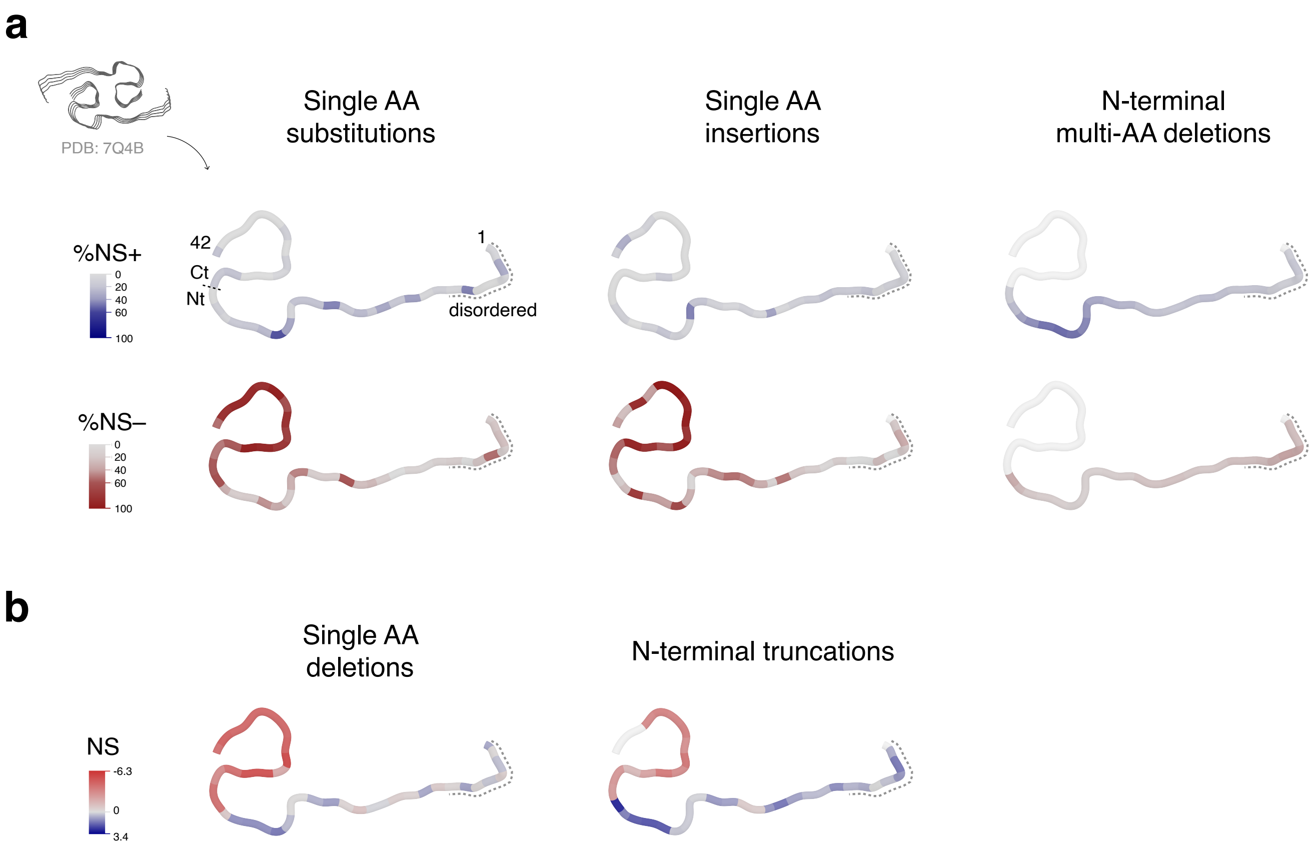
